# A mean-field approach for modeling the propagation of perturbations in biochemical reaction networks

**DOI:** 10.1101/2021.01.26.428329

**Authors:** Michelle Przedborski, David Sharon, Steven Chan, Mohammad Kohandel

## Abstract

Often, the time evolution of a biochemical reaction network is crucial for determining the effects of combining multiple pharmaceuticals. Here we illustrate a mathematical framework for modeling the dominant temporal behaviour of a complicated molecular pathway or biochemical reaction network in response to an arbitrary perturbation, such as resulting from the administration of a therapeutic agent. The method enables the determination of the temporal evolution of a target protein as the perturbation propagates through its regulatory network. The mathematical approach is particularly useful when the experimental data that is available for characterizing or parameterizing the regulatory network is limited or incomplete. To illustrate the method, we consider the examples of the regulatory networks for the target proteins c-Myc and Chop, which play an important role in venetoclax resistance in acute myeloid leukemia. First we show how the networks that regulate each target protein can be reduced to a mean-field model by identifying the distinct effects that groups of proteins in the regulatory network have on the target protein. Then we show how limited protein-level data can be used to further simplify the mean-field model to pinpoint the dominant effects of the network perturbation on the target protein. This enables a further reduction in the number of parameters in the model. The result is an ordinary differential equation model that captures the temporal evolution of the expression of a target protein when one or more proteins in its regulatory network have been perturbed. Finally, we show how the dominant effects predicted by the mathematical model agree with RNA sequencing data for the regulatory proteins comprising the molecular network, despite the model not having a priori knowledge of this data. Thus, while the approach gives a simplified model for the expression of the target protein, it allows for the interpretation of the effects of the perturbation on the regulatory network itself. This method can be easily extended to sets of target proteins to model components of a larger systems biology model, and provides an approach for partially integrating RNA sequencing data and protein expression data. Moreover, it is a general approach that can be used to study drug effects on specific protein(s) in any disease or condition.

## 1 Introduction

Systems biology has emerged in recent years as a powerful method for quantitative modeling of biological phenomena. The approach has gained attention because it is generic and it enables the integration of experimental data with mathematical and computational modeling to understand and make predictions about complex biological processes. Using a systems biology approach, one can model virtually any type of interacting system, including protein and gene networks, metabolic networks, cellular interactions, and tissue and organism-level interactions. The effectiveness of the method has contributed to its popularity for addressing questions in a number of different biological settings, including in molecular pathways underlying cancers and other diseases [1–13], in drug discovery and target identification [14–21], and in overcoming drug resistance [22, 23].

However, one limitation that arises in systems biology approaches, particularly in the context of molecular pathways and biochemical reaction networks, such as gene and protein regulatory networks, is that they often include several kinetic parameters that cannot be determined independently of experimental time series data. The limited availability of quantitative experimental data then leads to a trade-off in the development of the model. On one hand, the goal of a systems biology approach is to develop a mathematical model that is complex enough to capture the observed biological phenomena and to answer questions related to the corresponding protein or gene network. On the other hand, increasing the complexity of the model without sufficient experimental data leads to a model with poor predictive power [24]. In practice, fitting the outputs of the systems biology model to a limited experimental data set can lead to non-unique parameter sets or subsets of parameters that give fits of similar quality. This results in difficulties in interpreting the meanings of the parameters in the underlying molecular pathway or biochemical reaction network.

There are other methods for modeling the temporal evolution of gene regulatory networks, such as boolean models, that require only qualitative experimental data. However, while these types of models are easily extendable to large-scale systems, they tend to produce only qualitative results [24]. Another common issue that arises in the mathematical modeling of genetic pathways is that, in many cases, transcript levels alone are not sufficient to predict protein levels [25,26]. This is because RNA expression often does not correlate with protein expression for a number of reasons, including the availability of amino acids, the formation of higher order protein complexes, and post-translational modifications, such as phosphorylation and proteolytic events.

In this work we introduce a simplistic approach to model the temporal evolution of protein expression levels which partially overcomes the limitations and challenges faced by other modeling techniques. The method relies on the decomposition of a complicated molecular pathway into a mean-field model of the net effects exerted by the interaction network on a target protein. The method can be easily scaled to large interaction networks and can be generalized to encompass multiple target proteins and arbitrary types of perturbations to the network. Intriguingly, the dominant effects identified by the mean-field model, using only the target protein expression levels, are supported by RNA sequencing data of the regulatory proteins in the pathway, despite the model having no a priori knowledge of this data. Thus this approach offers interpretability to the modeling results and provides a stepping stone toward the integration of protein level data with gene expression data.

The manuscript is organized as follows. In Section 2, we discuss the relevant biological background and protein interactions for the examples considered in this work. In Section 3, we describe the experimental methods and the modeling approach, then present the results and discussion in Section 4. Finally, in Section 5 we give some concluding remarks.

## 2 Biology background

This work is motivated by the development of a large-scale mathematical model to study the effects of combination therapy on venetoclax-resistant acute myeloid leukemia (AML) cells [27,28]. A genome-wide CRISPR-Cas9 knockout screen revealed that genes involved in mitochondrial translation are potential therapeutic targets for restoring sensitivity to venetoclax in resistant AML cells [28]. The administration of antibiotics that inhibit mitochondrial protein synthesis, such as tedizolid and doxycycline, in combination with venetoclax, was observed to cause a heightened integrated stress response (ISR), resulting in potent cell death in resistant AML cells [28]. In addition to upregulating the ISR effector protein Chop, both drugs were observed to affect the expression of the c-Myc protein, which is known to play a significant role in hematopoiesis and whose overexpression is associated with lymphoma and leukemia [29].

At the center of the mathematical model describing the AML drug resistance pathway are the two transcription factors, c-Myc and Chop. Both transcription factors affect the expressions of the other proteins in the model, see Ref. [27]; however, they are themselves regulated by complex networks that are not included in the model. The mean-field approach was developed to quantitatively capture the time-dependent expression of the transcription factors under different experimental conditions, since their expression directly affects all other protein levels in the model. To this end, we first describe the regulatory networks that govern the expression levels of c-Myc and Chop, illustrating the complexity of these networks and highlighting the need for an approach that averages over interactions that are beyond the scope of the model.

### 2.1 The c-Myc oncogene

The c-Myc transcription factor is a protein that is encoded by the MYC oncogene and it is overexpressed and/or activated in more than 50% of human cancers [30]. The c-Myc protein plays a central role in apoptosis and cell growth and differentiation pathways, and its overexpression promotes tumorigenesis by activating key genes involved in glucose and glutamine metabolism, ribosomal and mitochondrial biogenesis, lipid synthesis, and cell-cycle progression [30–34]. Under normal conditions, proper levels of c-Myc are maintained throughout the cell cycle via phosphorylation and dephosphorylation events that tightly control its protein stability [31]. However, even under malignant conditions, the expression of c-Myc is affected by several pathways that control its rate of degradation and production, see Table 1. The c-Myc protein normally degrades quickly, with a half-life of 20-30 minutes [58, 59]. However, post-translational modifications, such as certain phosphorylation events, have been observed to increase the protein half-life up to as high as 80 minutes in some leukemia and lymphoma cell lines [58, 60] and as high as 178 minutes in non-small cell lung cancer cell lines in the presence of PIM2 overexpression [46]. Besides transcriptional regulation, the rate of production of c-Myc protein is also affected via translational control, and as we see in Table 1, some regulatory proteins promote cap-independent translation of c-Myc. In the case of 4E-BP, it is not clear whether the inhibition of cap-dependent translation and resulting promotion of cap-independent translation results in an increased or decreased rate of protein translation. However, internal initiation directed by the c-Myc internal ribosome entry site (IRES) was previously observed to be three-fold less efficient than cap-dependent translation initiation [56].

**Table 1:** Pathways that regulate the expression and stability of the transcription factor c-Myc. Unless otherwise specified, all residues and domains mentioned in the “mechanism” column refer to those belonging to the c-Myc protein.

| Pathway | Effect | Mechanism(s) | Reference |
| --- | --- | --- | --- |
| RAF/MEK1/2/<br>ERK1/2 | Increased protein stability | Phosphorylation at Ser62 | [31] |
| PI3K/AKT | Increased protein stability | Inhibits GSK-3β, preventing phosphorylation at T58 | [31] |
| USP28 | Increased protein stability | Deubiquitination | [35] |
| GSK-3β | Decreased protein stability | Phosphorylation at Thr58 (after phosphorylation at Ser62 by ERK) | [31] |
| UBB | Decreased protein stability | Degradation by the proteasome | [31] |
| JAK/STAT<br>(STAT3) | Increased production rate | Transcriptional activation | [36] |
| E2F1 | Increased production rate | Transcriptional activation | [37] |
| CEBPB | Increased production rate | Transcriptional activation | [38] |
| MEK2/3/MEK5/<br>ERK5 | Increased production rate | Transcriptional activation | [39] |
| p38 MAPK | Increased production rate | Transcriptional activation | [40] |
| NOTCH1 | Increased production rate | Transcriptional activation | [41] |
| TCF7L1/TCF7L2/<br>LEF1:CTNNB1 complexes | Increased production rate | Transcriptional activation | [42–44] |
| SIRT1 | Increased protein stability;<br>increased production rate | Deacetylation and linkage,<br>phosphorylation at Ser62, others<br>(double positive feedback loop) | [45] |
| FLT3-ITD/STAT5/<br>PIM1/2 | Increased production rate<br>via STAT5; increased protein<br>stability via PIM1/2 | Transcriptional activation;<br>phosphorylation at Ser329 by PIM2;<br>phosphorylation at Ser62 by PIM1 | [46, 47] |
| CEBPA | Decreased production rate | Transcriptional repression | [48] |
| p53 | Decreased production rate | Transcriptional repression | [32, 49, 50] |
| STAT1 | Decreased production rate | Transcriptional repression<br>(double negative feedback loop) | [51, 52] |
| RBL1:E2F4/5:DP1/2:<br>SMAD2/3/4 complex | Decreased production rate | Transcriptional repression | [53] |
| RUNX3 | Decreased production rate | Transcriptional repression due to<br>association with TCF7L1/2/LEF1:<br>CTNNB1 complexes | [54] |
| 4E-BP | Rate of protein translation | Binds to EIF4E, inhibiting cap-dependent translation and promoting<br>cap-independent translation by<br>freeing up ribosomes | [55, 56] |
| EIF4G2 (DAP5) | Rate of protein translation | Promotes IRES-mediated<br>cap-independent translation | [55, 56] |
| MIR34B/C | Decreased protein translation | Reduces the stability of c-Myc mRNA | [57] |

Not knowing precisely how the regulatory mechanisms affect the expression of the c-Myc protein is one challenge in developing a model of its regulatory network. Additional complexities arise due to the number of regulatory proteins, the feedback loops between c-Myc and the regulatory proteins, and the fact that several regulatory mechanisms depend on the phosphorylation status of the regulatory protein and/or the c-Myc protein. To complicate the situation further, several of the regulatory proteins also interact with each other, for example, GSK-3*β* and PI3K/AKT. Thus, developing a comprehensive model that accounts for all of the regulatory interactions described in Table 1 would require detailed protein expression data for each of the proteins and their phosphorylation status, which is typically not feasible due to experimental or cost limitations. In the absence of such detailed experimental measurements, a less complex approach must be used to capture the dynamics of the regulatory network.

### 2.2 Integrated stress response

The integrated stress response (ISR) is an intricate, evolutionarily conserved stress response pathway present in eukaryotic cells whose primary role is to restore cellular homeostasis. However, sustained or heightened activation of the ISR can lead to the expression of downstream effector proteins, such as Chop, resulting in the induction of apoptosis [61–63]. Depending on the cellular conditions, sustained ISR signalling can alternately cause cell death via ATF4-dependent necrosis or autophagic cell death [61].

The EIF2*α* kinases are the first responders to cellular stress signals, and the precise physiological and/or pathological conditions determine which of the kinases are activated. For example, ER stress can be caused by the accumulation of unfolded proteins in the ER, calcium homeostasis, perturbations in cellular energy, or redox status [61], resulting in the activation of PERK [62, 64–66]. In contrast, HEME deprivation leads to HRI activation and viral infection activates PKR, while amino acid deprivation, UV light, glucose depletion, and mitochondrial dysfunction lead to GCN2 activation [61, 64–67]. Some EIF2*α* kinases, such as PERK and GCN2, have signalling redundancy under certain stress conditions, and some cellular stressors, such as oxidative stress, can stimulate all of the EIF2*α* kinases [61].

The more specific stress responses, such as the mitochondrial unfolded protein response, UPR^MT^, or the unfolded protein response in the endoplasmic reticulum, UPR^ER^, are intimately connected with the more general ISR and consequently converge on a common signalling pathway. Specifically, stimulation of one or more of the EIF2*α* kinases leads to the phosphorylation of EIF2*α*, subsequently resulting in the attenuation of global protein synthesis. During this process, proteins with short upstream open reading frames (uORFs) in their 5’ untranslated region (5’UTR), such as the transcription factor ATF4, are preferentially translated to aid in cell survival and recovery [61]. The ATF4 protein is also regulated at the transcriptional and post-translational level, and its transcriptional activity and interactions with other transcription factors enable the ISR to produce distinct tailored responses to different cellular stress conditions [61].

The transcription factor ATF4 induces the transcription of several stress response genes, including Chop and GADD34. The mRNA of Chop and GADD34 both contain a 5’ uORF and are preferentially translated during the stress response; however the precise mechanisms controlling their translation are unclear [63]. While GADD34 forms a negative feedback loop by dephosphorylating EIF2*α* and restoring global protein synthesis, it has also been shown to play a role in the induction of apoptosis [61,62]. In a delayed response to prolonged conditions of cellular stress, ATF4 also transcriptionally activates 4E-BP. This pathway is a recently proposed second node of global translational inhibition in the ISR that shifts the cellular response toward cap-independent translation [55], resulting in increased translation of Chop and other stress response genes [68, 69]. The second node of translational inhibition is also mediated by stress-induced changes in the expression of mTOR, affecting the phosphorylation status of 4E-BP as well as the stability of ATF4 mRNA and its rate of translation [70].

The interactions described in this section shed light on the complexity of the ISR; however, there are a number of additional stress response genes and pathways that can be activated depending on the physiological conditions, many of which directly or indirectly affect the expression of Chop, see Table 2. As Table 2 indicates, in addition to transcriptional and translational regulation, Chop is regulated by post-translational modifications that affect its protein stability, which has been shown to have a half-life of four hours or less [71]. While the intricacy of the molecular pathway that regulates the expression of Chop can be inferred from Table 2, the complexity of the network is further compounded by additional interactions that occur between the regulatory pathways and the individual protein species.

**Table 2:** Pathways affecting the regulation of transcription factor Chop. Unless otherwise specified, all residues and domains mentioned in the “mechanism” column refer to those on the Chop protein. ISR-integrated stress response, UPR^ER^-unfolded protein response in the endoplasmic reticulum, UPR^MT^-unfolded protein response in the mitochondria. “p-” indicates phosphorylation.

| Pathway | Effect | Mechanism | Reference |
| --- | --- | --- | --- |
| PERK/p-EIF2α/ATF4 | Increased production rate | Transcriptional activation (ISR; UPR<sup>ER</sup>) | [61, 64–66]; [62] |
| GCN2/p-EIF2α/ATF4 | Increased production rate | Transcriptional activation (ISR; UPR<sup>MT</sup>) | [61, 64–66]; [67] |
| PKR/p-EIF2α/ATF4 | Increased production rate | Transcriptional activation (ISR) | [61, 64–66] |
| HRI/p-EIF2α/ATF4 | Increased production rate | Transcriptional activation (ISR) | [61, 64–66] |
| ATF6/ATF and cAMP, ATF6/ATF and cAMP/XBP-1 | Increased production rate | Transcriptional activation (UPR<sup>ER</sup>) | [62, 65, 66] |
| IRE1/XBP-1, IRE1/ASK1/JNK; IRE1/ASK1/p38MAPK | Increased production rate; Increased activity | Transcriptional activation; Phospho-activation at Ser78 and Ser81 (UPR<sup>ER</sup>) | [62, 65, 66]; [72–74] |
| PKR/c-JUN, JNK2/c-JUN | Increased production rate | Transcriptional activation (UPR<sup>MT</sup>) | [67, 75] |
| p-EIF2α-Ser51 | Increased production rate | Stimulates uORF-driven translation (attenuates global translation) (ISR, UPR<sup>ER</sup>; UPR<sup>MT</sup>) | [55, 69, 76–78]; [67] |
| mTOR/4E-BP1 | Increased production rate | mTOR phospho-inactivates 4E-BP proteins. **Decreased** expression of mTOR during ISR and UPR<sup>ER</sup> leads to hypo-phosphorylated 4E-BP1, which inhibits cap-dependent translation and promotes cap-independent translation. This increases CHOP uORF-driven translation (see below). | [55, 69, 77, 79] |
| p38MAPK/MNK1/p-EIF4E-Ser209, mTOR/4E-BP1, p-EIF2α-Ser51 | Increased production rate | These interactions were seen to collectively stimulate uORF-driven translation of CHOP. Particularly, increase in p-EIF4E and p-EIF2α and decrease in p-4E-BP1 (ISR and ER stress). | [69, 77] |
| ATF4/4E-BP | Increased production rate | ATF4 induces transcription of 4E-BP proteins (ISR, ER stress) | [55] |
| GADD34 | Decreased production rate | Dephosphorylates EIF2α, decreasing ATF4 expression (ISR, UPR<sup>ER</sup>) | [61] |
| TRB3 | Decreased production rate; Increased protein stability | Negative feedback on ATF4, mechanism unknown; inhibits association of p300 via N-terminal α-helix | [80, 81] |
| p300 | Decreased protein stability | Promotes ubiquitination by interacting with N-terminal domain | [80] |
| CypB | Decreased protein stability | Cooperates with p300 to promote ubiquitination | [82] |
| UBB | Decreased protein stability | Degradation by the proteasome | [80, 83] |
| AMPKα1 | Decreased protein stability | Promotes phosphorylation at Ser30, mediating ubiquitination | [83] |
| AKT | Decreased protein expression | Unknown (ER stress) | [84] |

Furthermore, while some regulatory pathways in the ISR, such as the role of ATF4, have been well-characterized, many details remain to be determined, such as how the ISR orchestrates a specialized translational response for different EIF2*α* kinases and stressors. In addition, a number of other mechanisms of preferential protein translation have been identified in cells, but their potential involvement in the ISR is currently unknown. For example, DAP5 is activated by caspase cleavage in apoptotic cells and promotes cap-independent translation of a number of stress responsive and apoptotic genes and cell cycle regulators such as c-Myc, Bcl-2, CDK1, Apaf-1, XIAP and c-IAP1, which could potentially play a role in stress resistance [55]. A complete characterization and understanding of the ISR under different physiological and pathological conditions, as well as the role of the ISR in disease progression, immune response, and long-term memory formation [63], will require systematic analysis and detailed experimental studies involving functional genomic tools and therapeutic manipulations. In the absence of such information, a simplistic approach for modeling the main dynamics in the regulatory pathway for Chop is imperative, especially when detailed experimental data is unavailable for model parameterization.

## 3 Methods

We developed a mean-field approach to overcome some of the limitations of standard modeling methods in the face of complex molecular pathways and limited experimental data. The approach developed here captures quantitative experimental trends as well as enables the interpretation of the pathway dynamics in response to a network perturbation, which, in this work, is due to the administration of therapeutic agents. The details of the available experimental data and the modeling approach developed here are explained below.

### 3.1 Experimental data

The experimental data set used in this work is comprised of in vitro measurements of c-Myc and Chop expression profiles in a line of venetoclax resistant AML cells, molm-13 R2 [28], under different treatment conditions. The c-Myc protein levels in treated cells were measured using intracellular flow cytometry analysis as previously described using an Alexa Fluor 647 conjugated anti-MYC antibody (Cell Signaling Technology, Cat # 13871). Chop protein expression was estimated by Chop reporter expression [28] and Chop mRNA levels. Chop mRNA levels in treated cells were measured using quantitative RT-PCR as previously described [28] using validated primers (OriGene, Cat # HP207450).

Due to experimental limitations, the resulting data set does not provide quantitative measurements of the initial concentration of c-Myc and Chop in untreated cells. Rather, the expression of c-Myc, relative to its initial concentration, was measured at *t* = 24, 48, and 72 hours post treatment administration, while the relative expression of Chop was measured at *t* = 24, 48, 72, and 96 hours post treatment administration. The treatment conditions consist of: single dose venetoclax monotherapy (400 nM concentration), single dose tedizolid monotherapy (5 *μ*M concentration), and single dose venetoclax and tedizolid combination therapy (400 nM and 5 *μ*M concentration, respectively), with the drugs administered simultaneously. No drug washout was performed during experiments, and the drugs were not replenished over the four day window.

The data set does not contain protein or RNA expression levels at *t* = 24, 48, or 72 hours post treatment administration for the c-Myc or Chop regulatory pathways described in Tables 1 and 2. However, RNA sequencing data was collected at *t* = 96 hours after treatment administration under each treatment condition, providing a snapshot of the gene expression profiles for all the species comprising the regulatory pathways. In Section 4 we compare the predictions of the mean field model with the RNA sequencing data to provide context to the predicted dominant interactions underlying the network dynamics for both c-Myc and Chop under each treatment condition.

### 3.2 Mathematical model

In the mean-field model, we aim to capture the temporal evolution of the target proteins c-Myc and Chop resulting from perturbations to their regulatory networks due to drug administration. In parallel, we aim to understand the dominant effects of the drug perturbations by identifying which pathways contribute most to the changes in c-Myc and Chop expression levels. Importantly, we stress that this approach is not limited to drug interactions. Rather, any chemical or physical perturbation which alters the regulatory pathway of the target protein can be modeled using this approach. The first step is to devise a mathematical description of the applied perturbation. Since, in this work we consider the effects of drugs administered to the system, we first develop a model of drug uptake in the cell culture before proceeding to the mean-field model in Section 3.2.2.

#### 3.2.1 Modeling drug pharmacokinetics

The in vitro cellular uptake and decay of drug *D* = {*A,T*} for venetoclax or tedizolid, respectively, was modelled as a system of two coupled ordinary differential equations (ODEs):

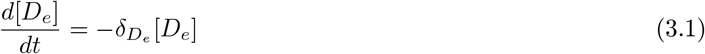

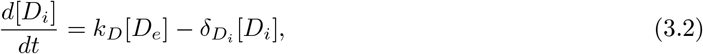

where *t* is the time, [*D_e_*] ≡ [*D_e_*(*t*)] denotes the extra-cellular concentration of drug *D*, and [*D_i_*] ≡ [*D_i_*(*t*)] is the intra-cellular concentration. This model assumes that the rate of drug uptake depends on the extra-cellular concentration and neglects depletion of the extra-cellular drug concentration due to drug uptake. Such a simplification is valid when the extra-cellular volume is much larger than the intracellular volume. Importantly, as discussed below, this simplification facilitates easily the incorporation of experimentally measured drug pharmacokinetics [85–88] into the model by suitable choices of the parameters *k_D_,δ_D_i__*, and *δ_D_e__*. Moreover, this choice of model allows for the investigation of the effects of changing the administered drug dose, which is useful for the process of optimizing the treatment protocol.

The typical time scale for experiments considering *in vitro* cellular drug uptake kinetics [86,89,90] is on the order of hours, which allows for the approximation [*D_e_*] ≈ const. = [*D*_0_], where [*D*_0_] is the initial administered drug concentration. Substituting this solution into Eq. (3.2) gives:

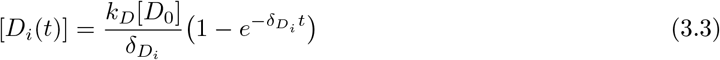

for the time evolution of the intra-cellular drug concentration. Solution (3.3) takes the value zero at *t* = 0 and saturates to the value *k_D_* [*D*_0_]/*δ_D_i__* when *t* ≫ 1/*δ_D_i__*, which is typically on the order of 30 minutes to several hours [86, 89, 90]. This solution qualitatively captures the cellular drug uptake curves that are observed in *in vitro* experiments [86, 89, 90], see, for example, Figure 1(a) for tedizolid *in vitro* drug uptake. Moreover, by a suitable choice of the parameters *k_D_* and *δ_D_i__*, Eq. (3.3) can provide quantitative agreement with short-term experimental *in vitro* drug uptake curves.

**Figure 1:**
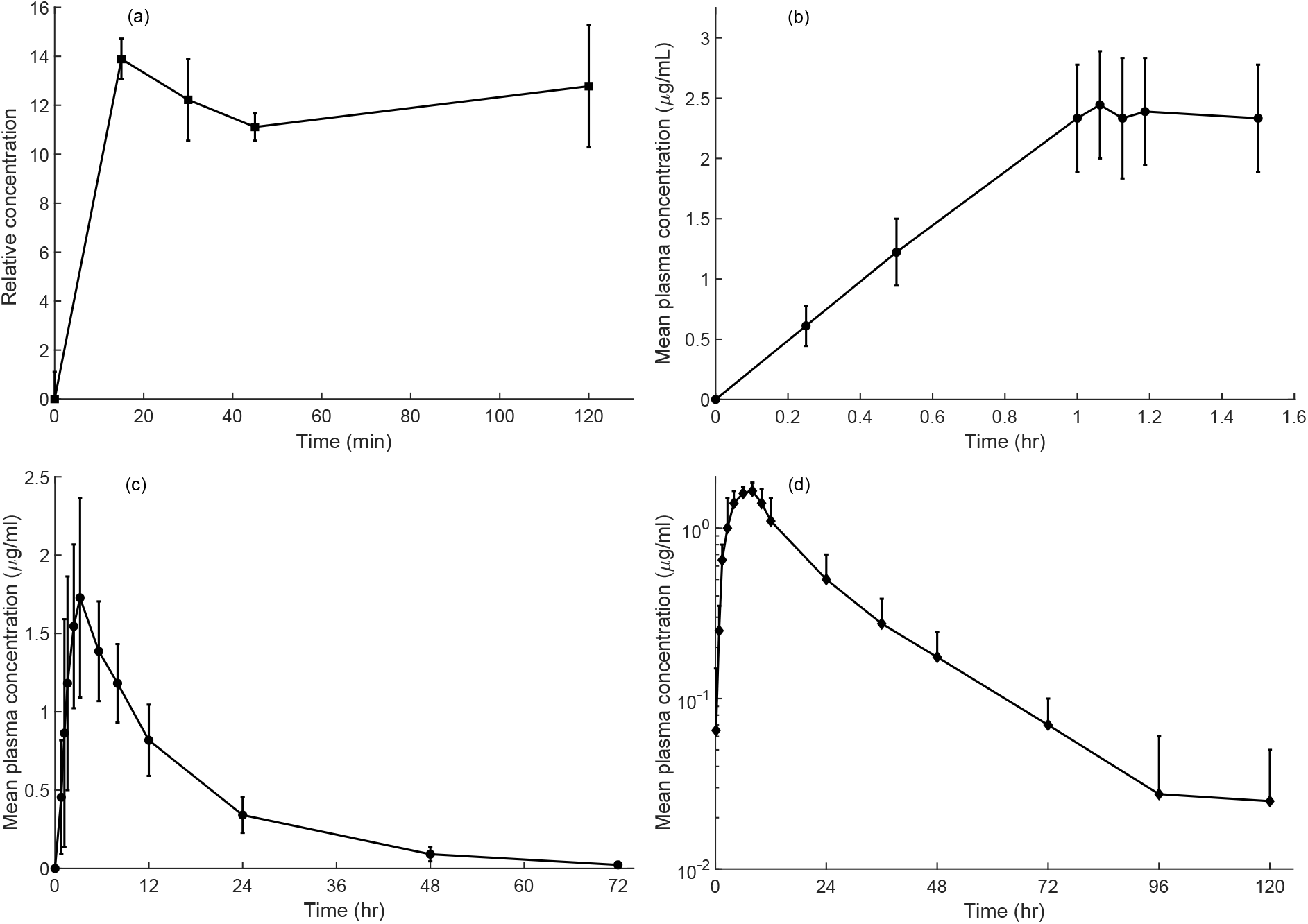
Experimentally measured cellular drug uptake curves under different environmental and drug administration conditions. (a) *In vitro* intra-cellular Tedizolid concentration (normalized to extra-cellular concentration) in THP-1 macrophage cells [86]. (b) Tedizolid *in vivo* drug plasma concentration, drug administered intravenously to patients [88]. (c) Long-term *in vivo* drug uptake curves for Tedizolid administered orally [88]. (d) Long-term *in vivo* drug uptake curve for venetoclax administered orally [91]. Plot shows total plasma concentration of venetoclax and its main metabolite M27. Images (a)-(d) were reproduced from Refs. [86], [88], and [91], and scales were chosen to facilitate direct comparison with the reported experimental data.

Interestingly, the drug uptake model given by Eqs. (3.1) and (3.2) can be used as a coarse-grained model for *in vivo* drug uptake by modifying the interpretation of [*D_i_*] to be the blood plasma concentration (assumed to be equal to the intra-cellular drug concentration), and [*D_e_*] to be the extra-vascular concentration. In this case, due to drug absorption, distribution, metabolism, and excretion from the body, [*D_e_*] cannot be taken to be constant for time scales *t* ≳ 1 hour, see for example, Figures 1(b) and (c). Rather, [*D_e_*] decays approximately exponentially, see for example Figures 1(c) and (d), and we incorporate the experimentally-determined apparent drug half-life t_1/2_ into the drug decay rate in Eq. (3.1) by setting

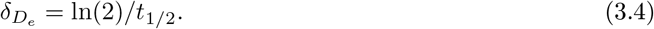

To choose appropriate values for the parameters *k_D_* and *δ_D_i__* that lead to the experimentally observed *in vivo* drug uptake curves, we begin by substituting the solution to Eq. (3.1),

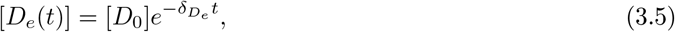

where [*D*_0_] is the initial concentration of drug administered, into Eq. (3.2) to obtain

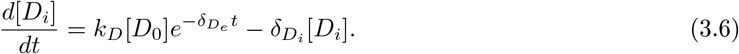

This is a separable ordinary differential equation (ODE) for [*D_i_*(*t*)], with solution

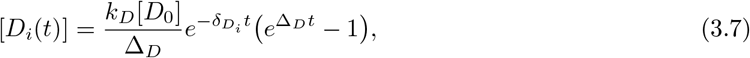

where

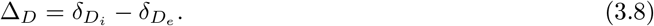

The solution in Eq. (3.7) takes on the value zero at *t* = 0 and has a peak at *t* = *t\**, after which it decays back to zero as *t* → ∞. To obtain the position of the peak *t*\*, we substitute the solution (3.7) into the right hand side of Eq. (3.6) and take 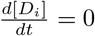, which gives:

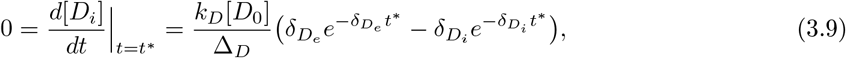

after simplification. Solving this last equation for *t\** gives

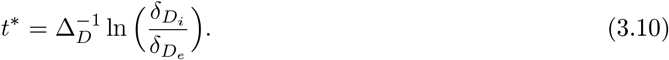

The peak intra-cellular drug concentration is next obtained by substituting *t* = *t*\* into Eq. (3.7), which gives

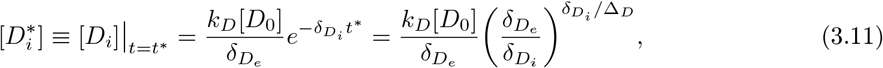

where the last equality follows from substitution of Eq. (3.10) into the expression. Next we express the peak intra-cellular drug concentration as 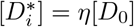, where 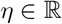, then we can isolate *k_D_* in Eq. (3.11), giving:

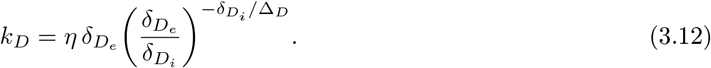

Equating *t\**, Eq. (3.10), to the experimentally-determined time to peak drug concentration gives a nonlinear equation for *δ_D_i__* in terms of *δ_D_e__* and *t\**, which can be solved numerically. The constant *η* can be determined from the experimentally-measured peak intra-cellular drug concentration, and then *k_D_* can be subsequently calculated from Eq. (3.12). Solving for the parameters *δ_D_i__* and *k_D_* in this way ensures that the simulated cellular drug uptake curve captures the experimentally measured *in vivo* drug pharmacokinetics, which are presented in Table 3 for venetoclax and tedizolid.

**Table 3:** Experimentally measured average *in vivo* drug pharmacokinetics obtained from Refs. [85–88] for venetoclax and tedizolid.

| Drug | Half-life<br>$\tau_{1/2}$ (hour<sup>-1</sup>) | Time to peak<br>concentration $t^*$ (hour) |
| --- | --- | --- |
| venetoclax | 23 | 5 |
| tedizolid | 10 | 2.5 |

For long-term (*t* ≳ 24 hours) *in vitro* experiments without drug washout or drug replenishment, the approximation [*D_e_*] ≈ const. = [*D*_0_] may not be appropriate due to the natural decay of the drug. In this case, a similar approach to the one described for *in vivo* drug uptake can be used to approximate the long-term *in vitro* cellular drug uptake, where the decay rate *δ_De_* is expected to be smaller than the corresponding *in vivo* parameter value.

In the next subsection we discuss how the mathematical model for drug pharmacokinetics developed here is integrated with the mathematical model for transcription factor regulation.

#### 3.2.2 Mean-field approach for modeling perturbations to biochemical reaction networks

As indicated in Tables 1 and 2, the transcription factors c-Myc and Chop, hereafter referred to as the “target” proteins, are each regulated by complex networks of interacting proteins that modify their stability (half-lives) and/or their production rates. A perturbation to one or more of the regulatory proteins will propagate through the network, ultimately affecting the temporal expression of each target protein. While it is possible to capture the full network dynamics with a detailed mathematical model, parameterization of the model would require experimental measurements of the temporal expression of each of the molecular species in Tables 1 and 2. In the absence of such detailed experimental data, a more coarse-grained approach, referred to as the mean-field approach, is better suited for modeling the temporal evolution of the target protein expression. In the mean-field approach developed in this work, the cumulative (net) *effect* of *groups* of regulatory proteins on the target protein level is modeled, rather than each of the individual regulatory proteins, see Figure 2. The development of this approach was guided by qualitative changes in RNA and protein expression levels. While this approach does not capture the full extent of the underlying biological protein network, it has the advantage that it reduces the complexity of the mathematical model and minimizes the number of kinetic parameters, while still retaining sufficient detail to capture the dynamic expression of the target protein.

**Figure 2:**
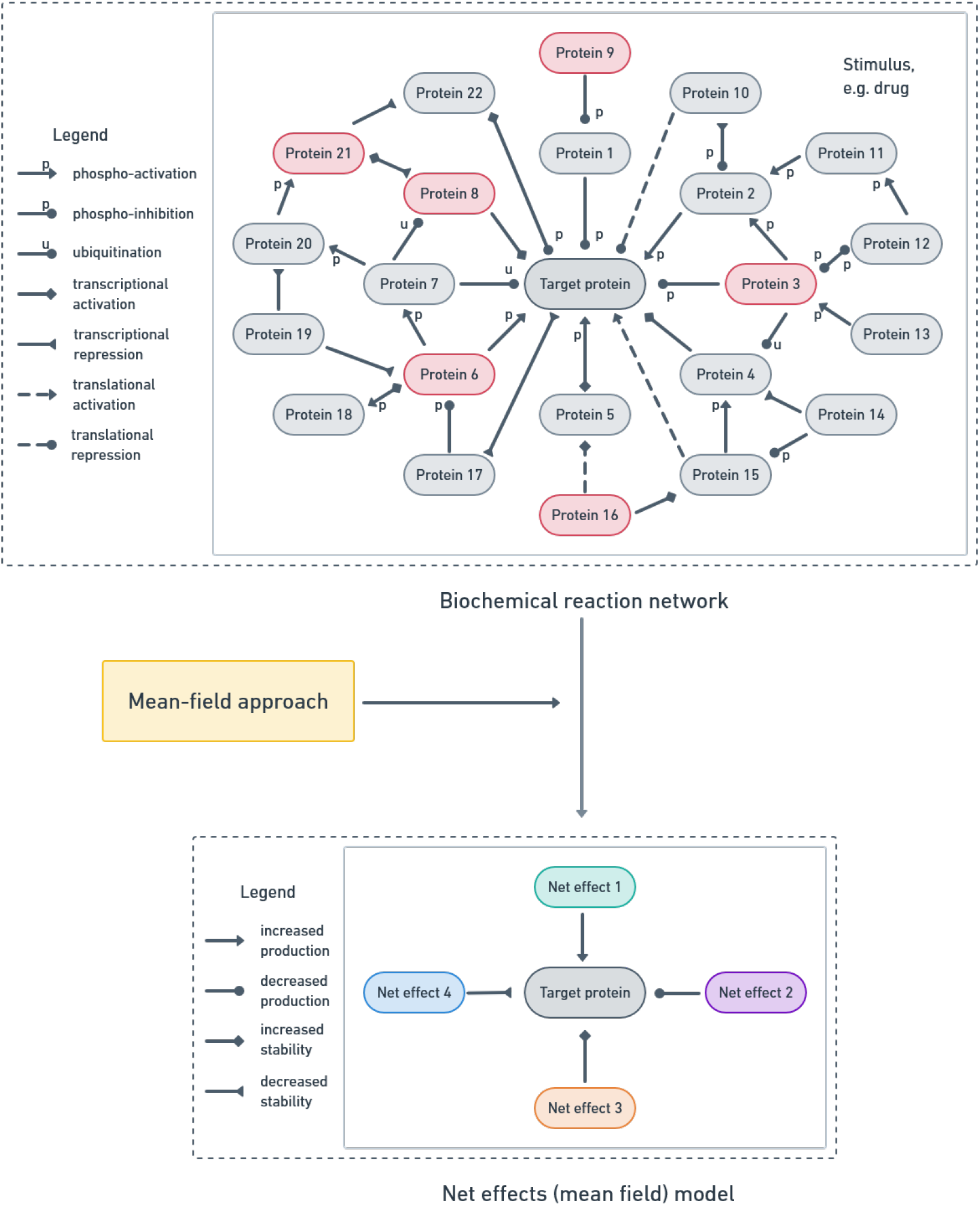
Depiction of the implementation of the mean-field approach for a biochemical regulatory reaction network. In the reaction network, the target protein refers to the chemical species whose temporal evolution we wish to model (in this work c-Myc or Chop), and the numbered proteins are regulatory proteins for the target protein, a fraction of which will be affected by drug administration. The result of the pipeline is a net effects model, wherein drug effects have a non-trivial temporal evolution that depends upon the drug concentration.

**Figure 3:**
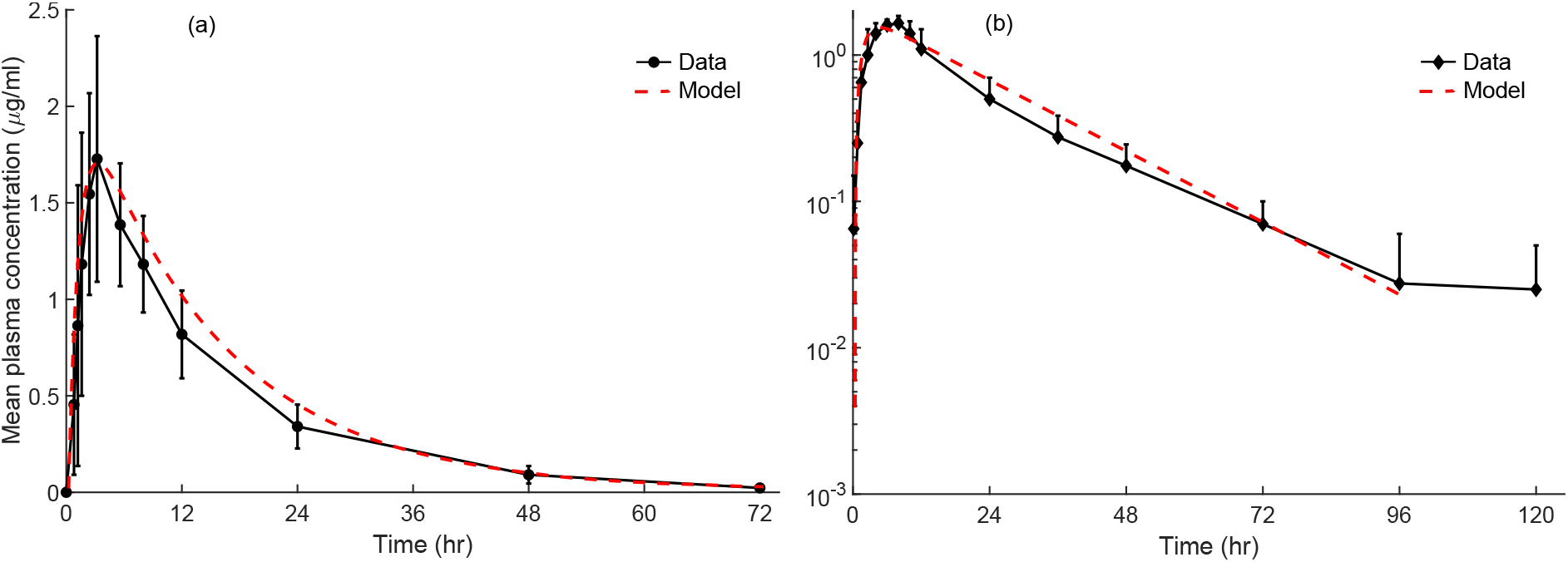
Simulated *in vivo* drug uptake curves corresponding to the parameter sets presented in Table 4. (a) Simulated *in vivo* drug uptake curve for tedizolid administered orally, shown as a dashed red line, captures the experimentally-measured plasma concentration over 72 hours [88]. (b) Simulated *in vivo* drug uptake curve for venetoclax administered orally, shown as a dashed red line, captures the experimentally-measured total plasma concentration (given by the total radioactivity curve) over 96 hours [91]. Experimentally-determined drug uptakes curves in (a), (b) were reproduced from Refs. [88], [91], and scales were chosen to facilitate direct comparison with the reported experimental data.

**Figure 4:**
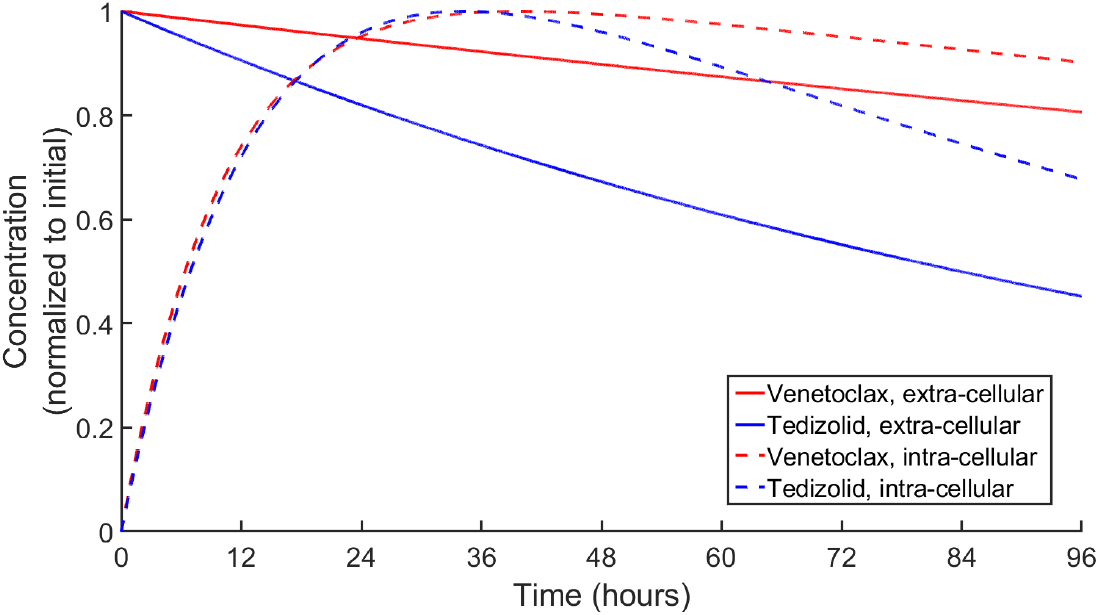
Simulated *in vitro* drug uptake curves corresponding to the parameter set presented in Table 5. Predicted long-term *in vitro* drug uptake curves for molm-13 R2 cells were determined by fitting to experimentally-measured c-Myc and Chop expression levels using the mean-field method.

In this work, we consider perturbations to the regulatory protein networks that are caused by the administration of therapeutic agents, see Section 3.2.1. To introduce the mean-field approach, we first consider the effects of a single therapeutic agent, i.e. venetoclax or tedizolid monotherapy. Given the regulatory pathways listed in Tables 1 and 2, we assume that monotherapy perturbs the regulatory network in a manner which effects the concentration of target protein, [*T_j_*], in four distinct ways: (1) increase in production rate, (2) decrease in production rate, (3) increase in stability, and (4) decrease in stability. The drug effects might occur at the protein level, for example, post-translational modifications of the protein product such as phosphorylation events; or the effects might occur at the RNA level, such as increased gene transcription, ultimately translating into protein-level changes. In the mean-field approach, we do not distinguish between the underlying mechanisms leading to each effect. Rather, an appropriate (mean) contribution is included in the model to capture each of the four drug effects on the temporal rate of change of [*T_j_*], resulting in the following first-order ODE:

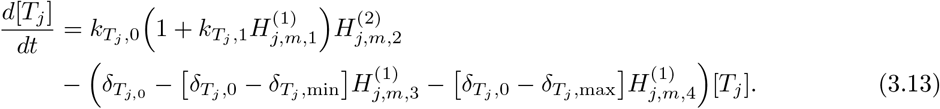

In the above equation, subscript *j* labels the target protein (*j* = 1, 2 corresponds to c-Myc or Chop, respectively), subscript *m* labels the drug (*m* = 1, 2 corresponds to venetoclax or tedizolid, respectively), *k*_*T_j_*,0_ is the background production rate for target protein *T_j_* in the untreated case, *k*_*T_j_*,1_ is a multiplier that dictates the maximum production rate for *T_j_* in the presence of drug, and *δ*_*T_j_*,0_, *δ*_*T_j_*,min_, and *δ*_*T_j_*,max_ are, respectively, the nominal, minimum and maximum decay rates for *T_j_*. We note that *k*_*T_j_*,0_, *k*_*T_j_*,1_, *δ*_*T_j_*,0_, *δ*_*T_j_*,min_, and *δ*_*T_j_*,max_ are biologically-constrained parameters that do not depend on the drug concentration. Rather, drug concentration is contained in the terms 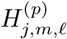 where *p* = 1, 2, which are Hill functions [92, 93] defined by:

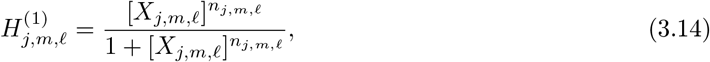

and

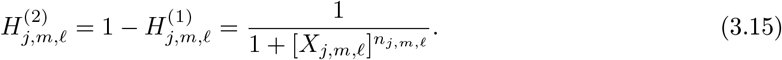

In Eqs. (3.14) and (3.15),

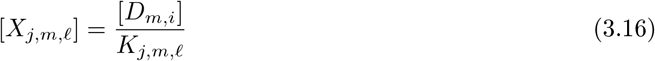

is the time-dependent intra-cellular drug concentration for drug *D_m_* (as indicated by the additional subscript “*i*” in [*D_m,i_*], see Subsection 3.2.1) normalized by the half-saturation constant *K_j,m,ℓ_* (where *K_j,m,ℓ_* > 0) for target protein *T_j_* and effect *ℓ*, where *ℓ* = 1,…, 4.

Hill functions of the form (3.14) take the value 0 at [*X_j,m,ℓ_*] ≪ 1 and saturate to the va [*X_j,m,ℓ_*] ≫ 1, which is consistent with stimulatory activity. In contrast, Hill functions of the form take the value 1 at [*X_j,m,ℓ_*] ≪ 1 and fall to 0 at [*X_j,m,ℓ_*] ≫ 1, representing inhibitory activity types of Hill functions satisfy 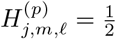 at [*X_j,m,ℓ_*] − 1 (i.e. [*D_m,i_*] = *K_j,m,ℓ_*). The shape and steepness of the Hill function is determined by the value of the exponent *n_j,m,ℓ_*, which is referred to the Hill as coefficient due to its origin in the context of ligand binding to macro-molecules [92]. In the context of protein expression, Hill functions are often used to model the transcriptional activation or transcriptional repression of the gene due to the binding of a transcription factor at the operator region(s) of the gene promoter [94]. We see from Eqs. (3.13)–(3.16) that in the mean-field approach, the temporal evolution of the target protein concentrations are modeled by treating the *drugs as transcriptional regulators*, assuming the quasi-steady state for mRNA expression levels [94]. This empirical approach enables the mathematical model to capture the main features exhibited by the experimental measurements of target protein expression, without requiring precise knowledge of each of the chemical species in the regulatory network, see Figure 2.

It can be easily verified that in the absence of drugs, 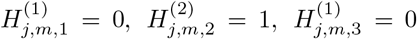, and 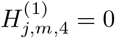, thus Eq. (3.13) reduces to:

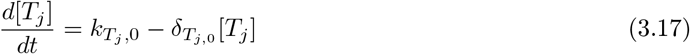

under control (untreated) conditions. From the above equation, it then becomes immediately clear that *k*_*T_j_*,0_ is indeed the background rate of production for [*T_j_*] in the absence of drugs, and *δ*_*T_j_*,0_ is the corresponding decay rate. Comparison of Eqs. (3.13) and (3.17) reveals that drug effect (1), increase in production rate, is captured by the term 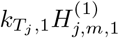 in Eq. (3.13), which increases the production rate to a maximum value of *k*_*T_j_*,0_(1 + *k*_*T_j_*,1_) when the drug concentration is sufficiently high. Drug effect (2), decrease in production rate, is captured by the term 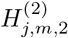, which reduces the total production rate by a factor of (1 + [*X_j,m,ℓ_*]*^n_j,m,ℓ_^*), allowing for the net production rate to fall below the background value if the drug concentration is sufficiently high. Drug effects (3) and (4), respectively, the increase and decrease in protein stability, are included in the model by incorporating two terms that perturb the decay rate of [*T_j_*] away from its nominal value *δ*_*T_j_*,0_. Specifically, drug effect (3) is captured in Eq. (3.13) by the term 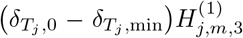, which gradually lowers the decay rate of [*T_j_*] to *δ*_*T_j_*,min_ as 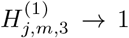. Similarly, drug effect (4) is captured by the term 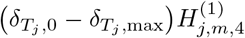, which gradually increases the decay rate to *δ_T_j_,max_* as 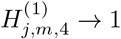. Both effects are maximal when the drug concentration is sufficiently high, and the nominal decay rate *δ*_*T_j_*,0_ is assumed to be constrained by *δ*_*T_j_*,0_ ∈ [*δ_T_j_,min_,δ_T_j_,max_*]. Similar to the drug effects (1) and (2), drug effects (3) and (4) are mutually competitive. In the event that both 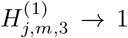 and 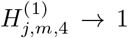, we see from Eq. (3.13) that the mean-field decay rate reduces to *δ_T_j__* = *δ*_*T_j_*,0_ + (*ϵ_max_* − *ϵ_min_*) where *ϵ_min_* = *δ*_*T_j_*,0_ − *δ_T_j_,min_* ≥ 0 and *ϵ_max_* = *δ_T_j_,max_* − *δ*_*T_j_*,0_ ≥ 0. We see that if *ϵ_min_* > *ϵ_max_*, drug effect (3) is stronger than drug effect (4) and *δ_T_j__* < *δ*_*T_j_*,0_, i.e. the protein is more stable. On the other hand, if *ϵ_max_* > *ϵ_min_*, drug effect (4) is stronger than drug effect (3) and *δ_T_j__* > *δ*_*T_j_*,0_, i.e. the protein is less stable.

Up to now, we have outlined a tractable framework for modeling the temporal evolution of a target protein whose regulatory network has been perturbed by a single therapeutic agent. Now we discuss how the framework can be extended to model the effects of combination therapy consisting of two or more therapeutics. Modeling the combined drug effects is not as simple as taking a linear superposition of individual drug effects for several reasons. First, due to the nonlinear nature of each drug effect (i.e. the Hill function form) it is expected that the effects of combination treatment should also take a nonlinear form. Second, it is possible for competition or co-operation to arise between the drug effects due to, respectively, antagonistic effects of each drug or nonzero overlap in the regulatory proteins that are affected by each drug. These interactions lead to additional nonlinearities in the functional form of the drug effects. To account for these nonlinearities, we follow the work of Refs. [94, 95] for the modeling of protein synthesis in the presence of multiple transcription factors. We begin by introducing the functions 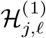 and 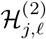 to describe the total drug effects resulting from the combination therapy, where *ℓ* =1, …, 4 as previously described, defined by:

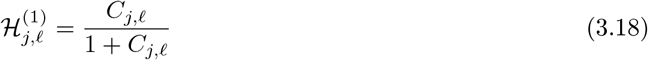

and

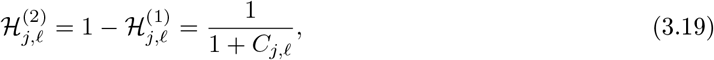

where

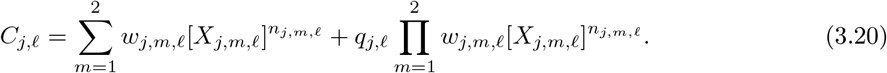

We see from the above equations that 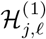 and 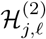 are Hill functions of the argument *C_jℓ_*, which as Eq. (3.20) illustrates, contains all instances of the drug concentrations, [*X_j,m,ℓ_*]. Taking *w_j,m,ℓ_* = 1 in Eq. (3.20) gives the familiar expression for protein regulation by two transcription factors, where cooperativity between transcription factor binding is described by the constant factor *q_j,ℓ_* [94, 95]. The case *q_j,ℓ_* = 1 corresponds to no cooperativity, *q_j,ℓ_* > 1 corresponds to positive cooperativity, and *q_j,ℓ_* < 1 to negative cooperativity.

In the context of drug effects, cooperativity between two or more drugs might arise when the protein networks affected by each drug overlap or have synergistic physiological functions. We follow the work of Ref. [95] to incorporate antagonistic drug effects, introducing the weights *w_j,m,ℓ_* in Eq. (3.20), where *w_j,m,ℓ_* ≥ 0 ∀ {*j,m,ℓ*}. The weights distinguish the relative importance of the individual drug effect to the combined effect during combination therapy. Taking *w_j,m,ℓ_* = 1 corresponds to each drug contributing equally to the combined effect, and this is taken to be the default value. Importantly, by taking *w_j,m,ℓ_* ≈ 0, the model captures cases where combined antagonistic drug effects nullify or dampen a pathway that plays a key role in the individual response to one or both drugs.

Incorporating all instances of the drug concentrations into Eq. (3.20) enables the mathematical model describing combination therapy to retain the same structure as the single drug treatment, Eq. (3.13). The previously described drug effects are captured in an identical way, but with the appropriate extension to the Hill functions 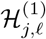 and 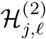. This gives:

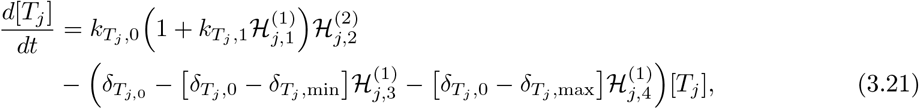

for the time evolution of target protein concentration [*T_j_*] during combination therapy. Importantly, while this approach was developed to model the effects of two drugs in combination, Eq. (3.21) can be further extended to model the effects of more than two drugs by appropriately generalizing the argument of the Hill functions, *C_j,ℓ_* to incorporate each of the drugs. Building on Eq. (3.20), this can be accomplished by first taking the sum in Eq. (3.20) to run over the number of distinct drugs. Then the second term in Eq. (3.20) can be generalized by taking the sum of all such pair-wise (second-order) products. In principle, one could also include a sum over higher order interactions. Doing so, in the case of three drugs, for example, the first sum would be comprised of three first-order terms, the second sum of three second-order terms, and the final term would be a single triple product of drug concentrations.

#### 3.2.3 Numerical simulations

We simulated the *in vitro* treatment protocols described in Section 3.1 using MATLAB version R2018a [96] with the ode15s differential equation solver. The cellular drug concentrations were simulated using Eqs. (3.1)–(3.2), where drug decay rates were taken to be parameters whose values were restricted to be smaller than the corresponding *in vivo* decay rates, see Table 3. We used Eqs. (3.13)–(3.16) to simulate the time evolution of c-Myc and Chop expression during venetoclax monotherapy and tedizolid monotherapy. The target protein expression during venetoclax/tedizolid combination therapy was simulated using Eqs. (3.18)–(3.21). For each target protein, the control parameters *K*_*T_j_*,0_, *δ*_*T_j_*,0_, and *K*_*T_j_*,1_ were kept the same for all treatment conditions.

Equations were non-dimensionalized in the time and protein domains. For each treatment option, the initial target protein levels were set to the value *K*_*T_j_*,0_/*δ*_*T_j_*,0_ (taken to be 1), which are the quasi-steadystate levels for the untreated system, see Eq. (3.17). The half-lives of c-Myc and Chop were constrained to lie between the experimentally-measured ranges reported in Section 2. The remaining parameter values were estimated by simulating the treatment strategies and forcing the simulated target protein levels to match to the experimental data using the MATLAB genetic algorithm. The loss function was taken to be the average relative difference between simulated and experimentally-measured data, with the average taken over all experimental time points, for both target proteins. The initial ranges for the parameter values were estimated based on biological relevance. After several generations of the genetic algorithm, each comprised of 200 iterations (simulations), the values of several estimated parameters were found to gravitate toward either the lower or upper bound of the estimated range. In this case, the parameter search range was updated approximately every 100 generations (or more often as needed to ensure sufficiently fast convergence), while attempting to maintain biological relevance. For example, for the regulatory effects, a strict upper bound was imposed on the Hill coefficients that tended toward larger values, signifying the switch-like nature of the effect.

In the initial run of the genetic algorithm, the parameter values were chosen randomly from within their respective initial ranges. Monotherapy simulations were first conducted to estimate the values of the parameters that are not specific to the combination treatment. Once a sufficient match to the experimental data was obtained, the combination therapy was additionally simulated. During these simulations, the weights and co-operativity factors were estimated and all parameter values were finetuned to sufficiently match to the three experimental data sets. Importantly, to reduce the number of parameters in the model, all the weights and co-operativity parameters were initialized to the value one, and were permitted to deviate from this value only if a sufficient match to the experimental data could not otherwise be obtained. These initial choices correspond to assuming that individual drug effects contribute equally to the combination therapy and that there is no cooperativity between drugs.

While the mean-field approach was developed assuming there are four possible drug effects for the target protein, it is feasible that not all effects are equally relevant to the time evolution of the protein. We would expect the experimental data to be reflective of the most dominant effects, and for these effects to similarly be reflected in the numerical simulations. By adjusting the kinetic parameter ranges as described above while fitting to experimental data with the genetic algorithm, we can identify and eliminate drug effects that are less-dominant to the overall response. Specifically, the half-saturation constants of the less-dominant effects will tend to diverge to a large value over fitting iterations, nullifying the contribution of the effect to the temporal evolution of the target protein.

After determining a nominal parameter set using the MATLAB genetic algorithm, we performed identifiability analysis to examine whether the parameter values were unique. Beginning with the nominal parameter set, we randomly generated 50,000 sets of parameter values using Latin hypercube sampling, with the lower and upper bounds set by ±*p*% of the nominal parameter value, with *p* typically taken to be 2, 5, or 15. For some parameters, such as the target protein or drug decay rates, a biologically relevant lower(upper) bound was imposed if it fell within the sampling range for the parameter. For each parameter set, we simulated the three treatment protocols and calculated the associated average relative error with respect to the experimental data. When the average relative error was below a threshold cutoff, we considered the corresponding parameter set *acceptable*; otherwise it was deemed *unacceptable*. If acceptable parameter sets were found, we further examined their statistical properties.

To incorporate temporal delays into the drug effects, we note that in the mean-field model, Eqs. (3.13) and (3.21), the drug concentration enters the equation through the argument of the Hill functions (3.14) and (3.15), and their extensions (3.18) and (3.19). Recognizing that Eq. (3.16) depends on the drug concentration at a particular time t, we make the replacement [*X_j,m,ℓ_*](*t*) = [*D_m,i_*](*t* − *τ_j,m,ℓ_*)/*K_j,m,ℓ_*, where *τ_j,m,ℓ_* is the time delay. Numerical integration of the mean-field model with these temporal delays was accomplished using the MATLAB dde23 delay-differential equation solver.

## 4 Results

Using the integrative methods described in Section 3, we examined the ability of the mathematical model to capture meaningful trends in the experimental data set for *in vitro* experiments of molm-13 R2 cells [28] treated with venetoclax monotherapy, tedizolid monotherapy, and venetoclax/tedizolid combination therapy. First, we show that the model for cellular drug uptake indeed captures the previously observed *in vivo* drug uptake curves. Then we illustrate that the long-term *in vitro* drug uptake curve predicted by the mathematical model is consistent with changes in RNA expression induced by adaptive drug resistance to venetoclax. We use identifiability analysis to show that only a small range of parameter values are consistent with the experimental data for the target proteins under the different treatment conditions. Then we illustrate how incorporating time delays between drug stimulus and biological response improves the match to the experimental data. Finally we show that the mean-field model can be interpreted as a data-driven approach by illustrating how predictions of the calibrated model agree with changes in RNA expression that are induced by treatment.

### 4.1 The mathematical model captures clinical drug pharmacokinetics

Using the method described in Section 3.2.1, we first sought to capture the drug uptake curves for orally-administered tedizolid and venetoclax in vivo. The resulting model parameter values are presented in Table 4, and the corresponding simulated intra-cellular (plasma) drug concentration curves are presented in Figures 1(a) and (b), along with the experimentally measured data for comparison. While the mean half-life of venetoclax was reported to be 23 hours in Ref. [91], the experimentally measured drug uptake curve in Figure 1(b) was more consistent with a half-life of ln(2) ×23 ≈ 16 hours. Nevertheless, comparison of the simulated *in vivo* drug curves with the experimentally-measured total drug concentrations in Figures 1(a) and (b) reveals that the simple system of two coupled ODEs, Eqs. (3.1) and (3.2), with appropriately chosen parameters, captures the main trends of *in vivo* drug uptake over a span of several days.

**Table 4:** Model drug parameter values corresponding to the experimentally-measured *in vivo* drug pharmacokinetics. In both cases, it was assumed that the peak intra-cellular concentration was the administered drug dose [*D*_0_] in the calculation of the parameter *k_D_*.

| Drug/condition | $\delta_{T_E}$<br>(hour<sup>-1</sup>) | $\delta_{T_i}$<br>(hour<sup>-1</sup>) | $k_D$<br>($[D_i]/[D_e]$·hour<sup>-1</sup>) |
| --- | --- | --- | --- |
| Venetoclax/*in vivo* | $\ln(2)/16 \approx 4.3 \times 10^{-2}$ | $\ln(2) \approx 6.9 \times 10^{-1}$ | $8.3 \times 10^{-1}$ |
| Tedizolid/*in vivo* | $\ln(2)/10 \approx 6.9 \times 10^{-2}$ | $4 \ln(2)/3 \approx 9.2 \times 10^{-1}$ | 1.1 |

**Table 5:**
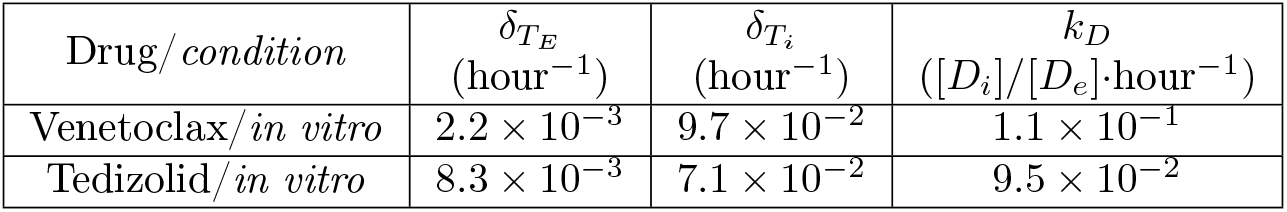
Optimal *in vitro* drug parameter values predicted by the mean-field model. In both cases, it was assumed that the peak intra-cellular concentration was the administered drug dose [*D*_0_] in the calculation of the parameter *k_D_*.

| Drug/condition | $\delta_{T_E}$<br>(hour<sup>-1</sup>) | $\delta_{T_i}$<br>(hour<sup>-1</sup>) | $k_D$<br>($[D_i]/[D_e]$·hour<sup>-1</sup>) |
| --- | --- | --- | --- |
| Venetoclax/*in vitro* | $2.2 \times 10^{-3}$ | $9.7 \times 10^{-2}$ | $1.1 \times 10^{-1}$ |
| Tedizolid/*in vitro* | $8.3 \times 10^{-3}$ | $7.1 \times 10^{-2}$ | $9.5 \times 10^{-2}$ |

### 4.2 Predicted slow *in vitro* cellular drug uptake is consistent with changes in RNA expression

The experimentally-measured target protein levels in molm-13 R2 cells were observed to change non-monotonically over several days during the *in vitro* treatment experiments. Importantly, the time-scale associated with the protein-level changes was consistent with non-constant cellular drug concentrations over the course of the experiment. This is because constant cellular drug concentrations would lead to a fixed change in the production and/or decay rates of the target proteins in the mathematical model, see Section 3.2.2. These fixed changes would cause the protein expression levels to reach a new steady state value on the timescale of the half-life of the proteins [94], which is at most on the order of hours.

Indeed, initial attempts to fit the experimental data for the target protein expression levels under the assumption of constant extra-cellular drug concentration were not successful at capturing the experimental trends due to this mismatch in timescales. Consequently, we enabled non-zero extra-cellular drug decay in the numerical simulations, where the decay rate was treated as a parameter whose value was constrained to be smaller than the reported *in vivo* drug decay rate, see Table 4. While some reports have indicated that the intra-cellular tedizolid concentration can reach up to 10-15 times the initial administered dose in *in vitro* settings [86], we made the simplifying assumption that the peak intra-cellular drug concentration is equal to the administered dose, i.e. *η* =1 for both tedizolid and venetoclax.

Incorporating non-zero extra-cellular *in vitro* drug decay enabled a good fit to the molm-13 R2 target protein data under all treatment protocols for the mean-field model. The predicted cellular drug uptake parameters are given in Table 4, and the corresponding simulated *in vitro* drug uptake curves are presented in Figure 1. A slightly different curve was obtained by including a temporal delay between drug stimulus and the biological response. However, for both drugs, the mathematical model predicted a slow rate of drug uptake, with the peak concentrations being reached at approximately 36 hours after drug administration, which is several times the reported time to peak drug concentration *in vivo* and in macrophages *in vitro*, see Table 3 and Figure 1.

This slow rate of drug uptake may be an artifact of the simplicity of the mathematical model. Further detailed experimental studies would be required to test the predictions of the drug uptake model. Nevertheless, there may be a biological basis for the predicted slow drug uptake by molm-13 R2 cells, even if the predicted drug uptake curves are not quantitatively accurate. Indeed, several mechanisms of drug resistance are known to prevent intra-cellular drug accumulation [97], including reduced drug influx due to cellular membrane changes, multi-drug resistance protein (P-gp) efflux, and drug entrapment in intra-cellular vesicles. Interestingly, each of these drug-resistance mechanisms is associated with an increase in sphingolipids and cholesterol membrane content [97].

To investigate the plausibility and putative mechanism of slow drug uptake in molm-13 R2 cells, we analyzed the RNA sequencing data from the venetoclax-resistant and venetoclax-sensitive molm-13 cell lines using the software DEseq [98]. This enabled quantification of the differences in equilibrium (untreated) total RNA levels between the two cell types. The most statistically significant differentially expressed genes were taken as those with adjusted *p*-value < 0.01, giving 6722 genes. These 6722 genes were sorted by adjusted *p*-value in ascending order, and the ranked list was input into GOrilla [99,100] for gene ontology analysis. The top 20 GO components, sorted in descending order by statistical significance, are presented in Table 6, from which it is clear that the sensitive and resistant molm-13 cells differ by several changes to the cell membrane structure and the formation of intra-cellular and extra-cellular vesicles. These cellular changes are consistent with a reduced rate of drug uptake, as predicted by the mathematical model.

**Table 6:**
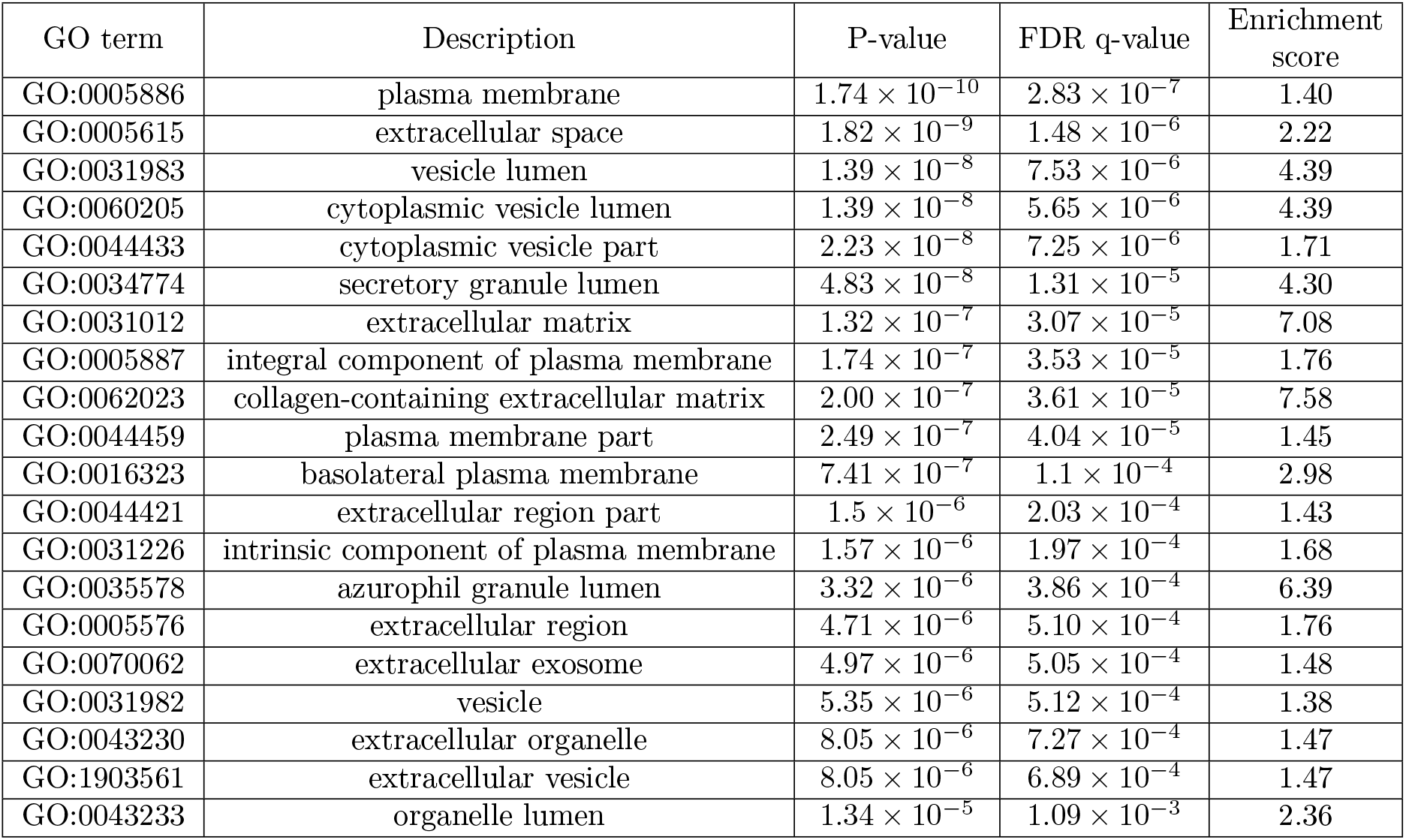
Top 20 enriched GO components obtained from analysis of RNA sequencing data from untreated venetoclax-resistant versus untreated venetoclax-sensitive molm-13 cells.

| GO term | Description | P-value | FDR q-value | Enrichment score |
| --- | --- | --- | --- | --- |
| GO:0005886 | plasma membrane | $1.74 \times 10^{-10}$ | $2.83 \times 10^{-7}$ | 1.40 |
| GO:0005615 | extracellular space | $1.82 \times 10^{-9}$ | $1.48 \times 10^{-6}$ | 2.22 |
| GO:0031983 | vesicle lumen | $1.39 \times 10^{-8}$ | $7.53 \times 10^{-6}$ | 4.39 |
| GO:0060205 | cytoplasmic vesicle lumen | $1.39 \times 10^{-8}$ | $5.65 \times 10^{-6}$ | 4.39 |
| GO:0044433 | cytoplasmic vesicle part | $2.23 \times 10^{-8}$ | $7.25 \times 10^{-6}$ | 1.71 |
| GO:0034774 | secretory granule lumen | $4.83 \times 10^{-8}$ | $1.31 \times 10^{-5}$ | 4.30 |
| GO:0031012 | extracellular matrix | $1.32 \times 10^{-7}$ | $3.07 \times 10^{-5}$ | 7.08 |
| GO:0005887 | integral component of plasma membrane | $1.74 \times 10^{-7}$ | $3.53 \times 10^{-5}$ | 1.76 |
| GO:0062023 | collagen-containing extracellular matrix | $2.00 \times 10^{-7}$ | $3.61 \times 10^{-5}$ | 7.58 |
| GO:0044459 | plasma membrane part | $2.49 \times 10^{-7}$ | $4.04 \times 10^{-5}$ | 1.45 |
| GO:0016323 | basolateral plasma membrane | $7.41 \times 10^{-7}$ | $1.1 \times 10^{-4}$ | 2.98 |
| GO:0044421 | extracellular region part | $1.5 \times 10^{-6}$ | $2.03 \times 10^{-4}$ | 1.43 |
| GO:0031226 | intrinsic component of plasma membrane | $1.57 \times 10^{-6}$ | $1.97 \times 10^{-4}$ | 1.68 |
| GO:0035578 | azurophil granule lumen | $3.32 \times 10^{-6}$ | $3.86 \times 10^{-4}$ | 6.39 |
| GO:0005576 | extracellular region | $4.71 \times 10^{-6}$ | $5.10 \times 10^{-4}$ | 1.76 |
| GO:0070062 | extracellular exosome | $4.97 \times 10^{-6}$ | $5.05 \times 10^{-4}$ | 1.48 |
| GO:0031982 | vesicle | $5.35 \times 10^{-6}$ | $5.12 \times 10^{-4}$ | 1.38 |
| GO:0043230 | extracellular organelle | $8.05 \times 10^{-6}$ | $7.27 \times 10^{-4}$ | 1.47 |
| GO:1903561 | extracellular vesicle | $8.05 \times 10^{-6}$ | $6.89 \times 10^{-4}$ | 1.47 |
| GO:0043233 | organelle lumen | $1.34 \times 10^{-5}$ | $1.09 \times 10^{-3}$ | 2.36 |

Furthermore, the RNA sequencing data indicated that molm-13 R2 cells exhibit a three-fold increase in ABCA1 RNA levels in comparison to the venetoclax-sensitive molm-13 cells (adjusted *p*-value 2.3×10^−100^). Along with the ABCB1 (multi-drug resistance 1) gene, which is well-known to play a substantial role in multi-drug resistance in several cancers, the protein encoded by the ABCA1 gene is a member of the superfamily of ATP-binding cassette (ABC) transporters. These are membrane-associated proteins that transport various molecules across extra- and intra-cellular membranes, and can expel drugs from the interior of the cell [101]. Indeed, increased ABCA1 expression has been previously observed to play a role in therapeutic resistance in other types of cancers [102, 103]. Thus, among other changes, the significant increase in ABCA1 expression observed in molm-13 R2 cells is consistent with the slow drug uptake predicted by the mathematical model.

### 4.3 The mean-field approach leads to narrow ranges for kinetic parameter values

Precisely half of the half-saturation constants drifted toward increasing values as the genetic algorithm converged on the experimental data for both target proteins. This indicated that the corresponding drug effects were not relevant to the temporal evolution of the target protein level, and consequently, these effects were removed from the model to reduce its complexity. Interestingly, this phenomenon, in which half of the drug effects automatically “drop out” of the model when fitting to the experimental data, occurred when the initial parameter values, treatment conditions, and ranges for the parameter values were varied. Thus, the integration of experimental data with the mean-field model can reveal the dominant drug effects on the target protein expression, enabling a reduction in the model complexity.

The simulated target protein levels that best fit to the experimental data are depicted in Figure 5, and the corresponding nominal parameter set is presented in Supplemental Table S1. The relative error was calculated between the output of the model and the average experimental measurement at each time point. The nominal parameter set led to an average relative error of 0.089 for both target proteins, over all time points and treatment conditions. For c-Myc, increased stability and decreased production were found to be the dominant drug effects for both venetoclax and tedizolid monotherapy, while for Chop it was decreased stability and increased production rate. Interestingly, positive cooperativity was predicted by the mean field model for the production rate effects of both target proteins during venetoclax and tedizolid combination therapy. Furthermore, the experimental data suggested that a deviation from *w_j,m,ℓ_* = 1 was necessary to capture the behaviour of c-Myc with combination treatment. This is because combination treatment led to a significant drop in c-Myc levels in molm-13 R2 cells, and the rebound back to the control value was slow, taking at least several days, see Figure 5. Taken together, this behaviour suggested that drug effects that increase c-Myc protein expression, while relevant for both monotherapies, are not dominant effects during the combination treatment.

**Figure 5:**
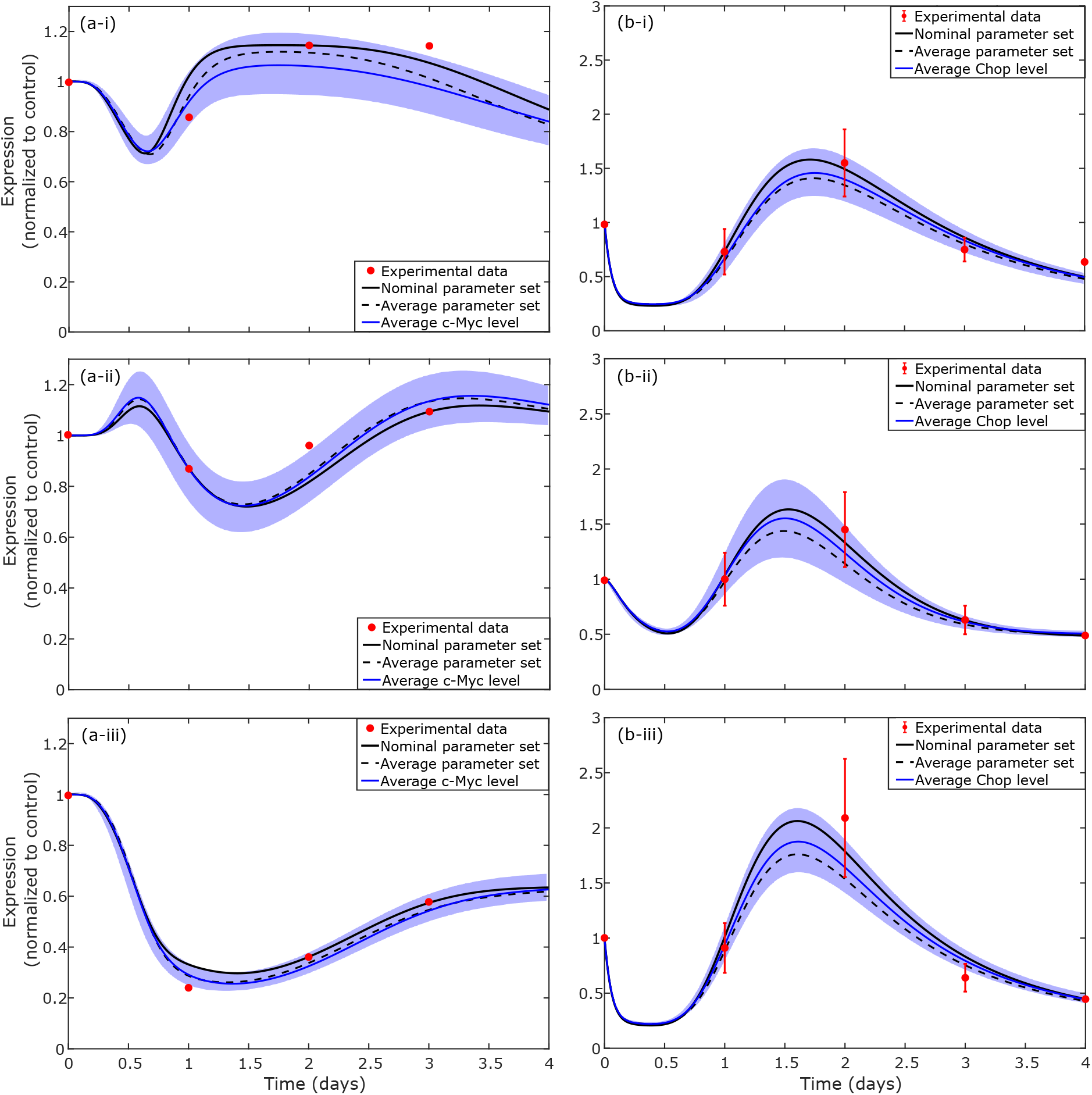
Target protein levels obtained from the mean-field model. Nominal parameter set refers to the best fit to the experimental data. Average parameter set refers to the averages calculated from identifiability analysis. Average target protein level indicates the average over the distributions produced by all parameter sets obtained from identifiability analysis. Standard deviation in target protein levels is indicated by the blue shaded area. Column (a) refers to c-Myc expression, column (b) to Chop expression. Row (i) refers to venetoclax monotherapy, (ii) to tedizolid monotherapy, and (iii) to venetoclax and tedizolid combination therapy.

We next explored the uniqueness of the parameter fit by performing identifiability analysis as described in Section 3.2.3. First we determined whether there were any parameter sets, other than the nominal parameter set, that lead to an average relative error of less than 0.09, which is slightly larger than the relative error associated with the nominal parameter set. When the ranges for Latin hypercube sampling (LHS) were set by deviations of ±5% of the nominal parameter values, only one of the 50,000 parameter sets sufficiently fit to the data. In comparison, when the sampling ranges for LHS were set by deviations of ±2% of the nominal parameter values, six of the 50,000 parameter sets fit the data with an average relative error of less than 0.09. The standard deviation of these six parameter sets was at most 1.3% of the parameter value, indicating that the acceptable parameter sets fall within a very narrow range of values.

We subsequently investigated the spread that arises in the target protein levels from increasing the error cutoff to a larger value. In this work, we chose 0.15, corresponding to an average relative error of up to 15% to be considered an acceptable fit to experimental data. In this case, 68 out of 50,000 sampled parameter sets fit to the data within the threshold error. The simulated target protein levels corresponding to the average parameter values, taken over all 68 sets, are depicted in Figure 5 in comparison with the nominal parameter set. The standard deviations in the parameter values in the 68 sets were at most approximately 9.5% of the parameter value. We also show in the Figure the average target protein levels at each time point along with their standard deviation, obtained from the distributions produced by the 68 parameter sets. Importantly, we see from Figure 5 that in some instances, the spread in the target protein is up to nearly 25% of the average protein level. These findings reinforce the observation that a close fit to the experimental data occurs only within a narrow range of parameter values.

### 4.4 Incorporating temporal delays into the drug effects better captures the experimental trends

We see from Figure 5 that the nominal parameter set in Supplemental Table S1 gives a fair match to the in vitro data, with an average error of less than 10% for all time points and treatment conditions, for both target proteins. Overall, the mean-field model introduced in Sec. 3.2.2 qualitatively captures the temporal evolution of the target protein levels. However, close inspection of Figure 5 suggests that the fit might be improved by incorporating a temporal delay into some of the drug effects, as described in Sec. 3.2.3. For example, for c-Myc with Tedizolid treatment, Figure 5 (a-ii), the model predicts a slight increase in c-Myc levels within the first 24 hours, and it does not quite capture the expression level on day 2. For Chop, the model predicts a significant decrease in expression within the first 24 hours for treatments containing venetoclax, and the predicted peak appears to occur slightly too early. Biologically speaking, temporal delays in the response to a drug might occur for several reasons. For example, consider a regulatory pathway for a target protein that is downstream of several other protein interactions. If the administration of a drug affects the expression of one of the proteins that is upstream of the regulatory pathway, there may be a series of interactions, including transcription and translation events and post-translational modifications, that must occur before the regulatory protein is affected. In this scenario, it seems reasonable to assume that there should be a time delay between drug administration and the corresponding effect on the target protein.

Indeed, incorporating temporal delays into the drug effects, see Sec. 3.2.3, reduced the average relative error by a factor of nearly three, to 0.0304, for all time points, treatment conditions, and both target proteins. Importantly, we note that additional iterations of the genetic algorithm would enable a further (albeit slow) decrease in the relative error and thus a better match to the experimental trends. The best fit to the experimental data is depicted in Figure 6 and the corresponding parameter set is presented in Supplemental Table S2. We see from Figure 6, that by incorporating temporal delays into the drug effects, the mean-field model can better capture both the quantitative and qualitative trends in the experimental data, c.f. Figures 5 and 6. Notably, an upregulation in c-Myc expression within the first 24 hours of tedizolid treatment is not predicted in the temporal delay model. Interestingly, the fit to the tail end of c-Myc expression during venetoclax treatment is also far superior in the temporal delay model. In addition, the mean-field model with temporal delay does not predict as significant of a decrease in Chop expression within the first 24 hours of venetoclax or combination treatment. As well, the peaks in Chop expression are slightly shifted forward in time to better match the experimental data in the model with temporal delay. We also verified the importance of including weights *w_j,m,ℓ_* = 1 for combination treatment in the case of the model with temporal delay. Forcing these weights to be equal to unity, the lowest average relative error we obtained was 0.13 for all time points, treatment conditions, and both target proteins.

**Figure 6:**
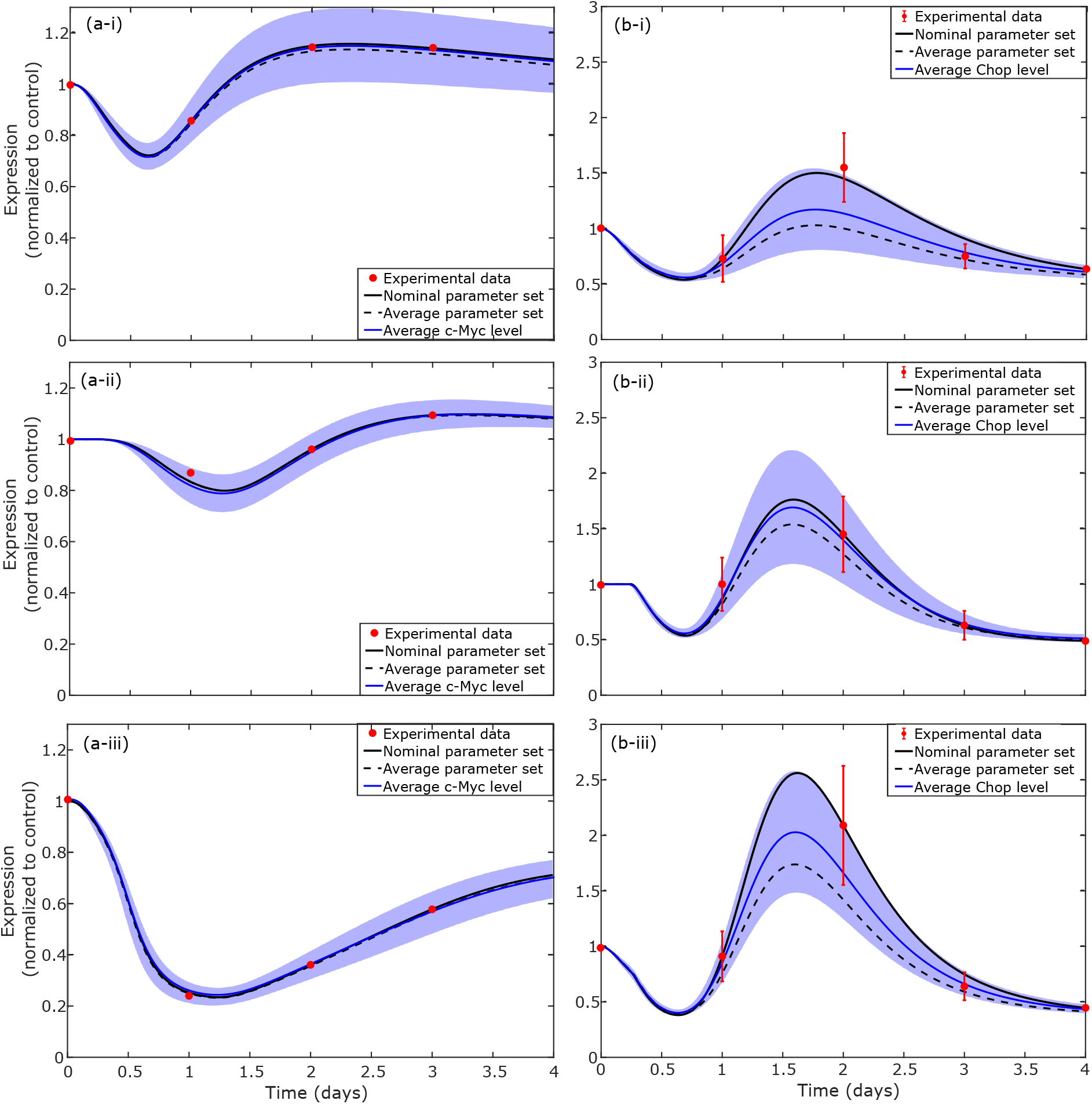
Target protein levels obtained from the mean-field model with temporal delays in the drug effects. Nominal parameter set refers to the best fit to the experimental data. Average parameter set refers to the averages calculated from identifiability analysis. Average target protein level indicates the average over the distributions produced by all parameter sets obtained from identifiability analysis. Standard deviation in target protein levels is indicated by the blue shaded area. Column (a) displays c-Myc expression, column (b) Chop expression. Row (i) depicts venetoclax monotherapy, (ii) tedizolid monotherapy, and (iii) venetoclax and tedizolid combination therapy.

We next explored the uniqueness of the parameter fit by performing identifiability analysis as described in Section 3.2.3. First we determined whether there were any parameter sets, other than the nominal parameter set, that lead to an average relative error of 0.0304 or less. When the ranges for Latin hypercube sampling (LHS) were set by deviations of ±5% of the nominal parameter values, none of the 50,000 parameter sets sufficiently fit to the data. Indeed, when the sampling ranges for LHS were set by deviations of ±2% or ±1% of the nominal parameter values, none of the 50,000 parameter sets fit the data with an average relative error of 0.0304 or less. Consequently, we set the deviation to an order of magnitude smaller, ±0.1%, and in this case we found that 386 of 50,000 parameter sets fit sufficiently well to the experimental data. The standard deviation of these 386 parameter sets was at most 0.06% of the parameter value, indicating that the acceptable parameter sets fall within an extremely narrow range of values for the mean-field model with temporal delay.

We subsequently investigated the spread that arises in the target protein levels from increasing the error cutoff to a larger value. We chose the cutoff to also be 0.15 for this model, corresponding to an average relative error of up to 15% to be considered an acceptable fit to experimental data. In this case, 3,744 out of 50,000 sampled parameter sets fit to the data within the threshold error. The simulated target protein levels corresponding to the average parameter values, taken over all the parameter sets, are depicted in Figure 6 in comparison with the nominal parameter set. The standard deviations in the parameter values in the 3,744 sets were at most approximately 8.8% of the parameter value. We also show in the Figure the average target protein levels at each time point along with their standard deviation, obtained from the distributions produced by the 3,744 parameter sets. Importantly, we see from Figure 6 that in some instances, the spread in the target protein is over 70% of the average level, despite the small changes in the parameter values. These findings reinforce the observation that a close fit to the experimental data occurs only within a narrow range of parameter values.

### 4.5 Dominant effects predicted by the mean-field approach are supported by differential gene expression

Lastly, we examined whether there was a correlation between the significant drug effects predicted by the mean-field model and the RNA expression levels under different treatment conditions. This analysis enabled insight into how drug effects at the protein level are reflected in the changes at the gene level. To this end, we performed differential gene expression analysis on the RNA sequencing data for each treatment condition using the software DEseq [98]. We then developed a test statistic *ζ* that includes both the magnitude of the change in RNA expression as well as the adjusted *p*-value to quantify the significance of the change. Since the adjusted p-values lie within the interval *p* ∈ [0,1], and thus log_2_(*p*) ≤ 0, for each gene *i* we define:

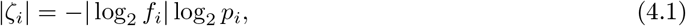

where *f_i_* is the fold-change of the normalized RNA-sequencing data (RNA-seqt_reated_/RNA-seq_control_) for the gene. To capture the nature of each drug effect on the target protein level, the sign of the test statistic *ζ_i_* is assigned as follows. If gene *i* is known to have a stimulatory effect on the target protein, such as increased production or increased stability, then sgn(*ζ_i_*) = sgn(log_2_(*f_i_*)); while if gene *i* is known to have an inhibitory effect, such as decreased production or decreased stability, then sgn(*ζ_i_*) = −sgn(log_2_(*f_i_*)). In this way, the test statistic directly captures the change in the target protein level resulting from changes in the regulatory genes. Specifically, an increase(decrease) in a stimulatory gene is taken to be positive(negative), while an increase(decrease) in an inhibitory gene is taken to be negative(positive).

Next, we analyzed the value of the test statistic *ζ* for the dominant regulatory genes in Tables 1 and 2. This enables the identification of the drug effects that are most significant to the expression of the target protein. The results are depicted in Figure 7 for both c-Myc and Chop. We see from Figure 7(a) that, overall, there is a net increase in the production rate for c-Myc under all three treatment conditions, as well decreased protein stability, as predicted by the mean-field model with temporal delay. During tedizolid monotherapy and venetoclax/tedizolid combination therapy, several pathways that affect the production rate of c-Myc, with both positive and negative regulatory effects, are perturbed. Importantly, the test statistic indicated that combination treatment leads to the smallest production rate for c-Myc, which supports the predictions of the mean-field model with temporal delay and the experimental trends in protein expression. Combination treatment is also seen to lead to the least stable c-Myc protein, which confirms additional predictions of the mean-field model with temporal delay.

We see from Figure 7(b) that the production rate for Chop also increases with each treatment condition, while the stability of Chop decreases, as predicted by the mean-field model (with and without temporal delay). In addition, both the rate of production and the instability in Chop were found to be maximal with combination treatment, which is reflected in the in vitro data and the predictions of the mean-field model (with and without temporal delay), see Figures 5 and 6. Interestingly, while both stimulatory and inhibitory effects are seen to be important for the production of Chop, Figure 7(b) is evidently far more sparse than Figure 7(a), indicating that the perturbed regulatory network for Chop is not as complex as the perturbed regulatory network for c-Myc. It should be noted that in Figure 7, genes that promote cap-independent protein translation of c-Myc were taken to decrease its production rate, relative to the control. This is because cap-independent translation initiation is well-known to be less efficient than the cap-dependent one [104].

**Figure 7:**
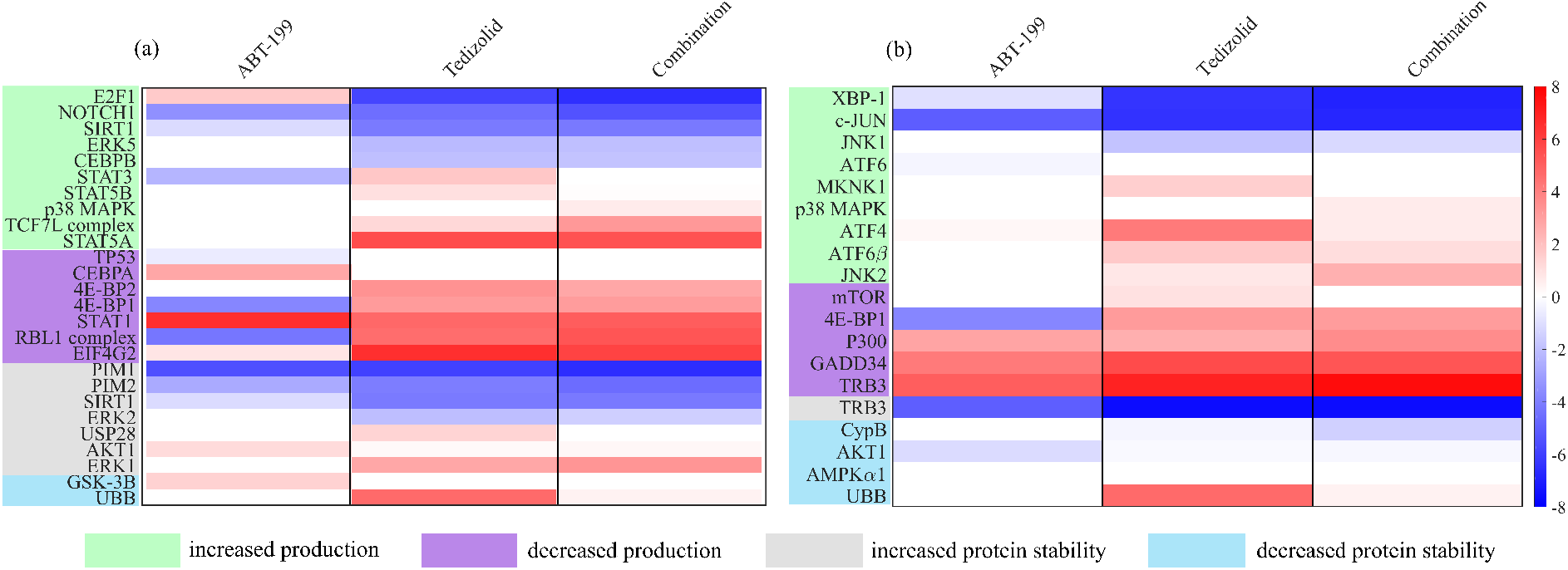
Effects of changes in the regulatory genes on the target proteins (a) c-Myc and (b) Chop,
obtained via differential gene expression analysis. Specifically, we plot sgn(*ζ_i_*) ln (abs(*ζ_i_*)), where the sign of *ζ_i_* depends on the type of regulatory effect, as explained in the text. Positive values (red bars) indicate a resulting increase in protein expression, while negative values (blue bars) indicate a decrease. For each drug effect, the genes are arranged in order of increasing value of sgn(*ζ_i_*) ln (abs(*ζ_i_*)) for combination
treatment.

## 5 Discussion

In this work, we constructed an analytical and computational framework for modeling the temporal propagation of perturbations through a regulatory gene network. The framework relies on first identifying the possible regulatory effects that can occur for a protein of interest, referred to in this work as the target protein. These effects are subsequently implemented into a mean-field mathematical model to capture the dominant regulatory behaviour leading to the time evolution of the target protein. To illustrate the method, we applied the approach to study the time evolution of the target proteins c-Myc and Chop during venetoclax and tedizolid monotherapy and venetoclax/tedizolid combination treatment in resistant molm-13 AML cells. Both target proteins have regulatory networks that are highly connected and comprised of several distinct species. We showed that the predictions of the model agree with the temporal evolution of protein data for c-Myc and Chop, as well as RNA sequencing data for the regulatory protein network. Importantly, the approach gives insight into the most significant interactions that are underlying the observed temporal response, which is particularly valuable when the temporal data for the regulatory network is limited.

This empirical approach is highly useful when the underlying regulatory network is comprised of many molecular species, has a high connectivity, and/or a significant fraction of the regulatory genes (proteins) are affected by the perturbations to the network. Interestingly, findings from a recent report [105] indicate that several of these network features are exhibited in aggressive B-cell lymphomas during venetoclax treatment and in the development of resistance to venetoclax. Specifically, in addition to its intended role as a Bcl-2 protein inhibitor, venetoclax was observed to result in the activation of several common genes in multiple mantle cell lymphoma and diffuse large B-cell lymphoma cell lines, including those involved in apoptosis, DNA damage response/repair, cell growth/survival signaling pathways, metabolism, and mitochondrial genes [105]. In these cell lines, a total of 15 proteins were found to be commonly significantly up-regulated (or activated via phosphorylation), including: p-SRC, p-MEK1, p-MAPK, p-JNK, p-AKT, GCLM, and TFAM, where “p-” indicates the phosphorylated state. Moreover, a total of 16 proteins were commonly significantly down-regulated, including: CHK2, mTOR, p38 MAPK, PTEN, RICTOR, RIP, and STAT5A. Significantly, comparison with Tables 1 and 2, indicate that several of these common differentially expressed proteins control the expression of the target proteins c-Myc and/or Chop, considered in this work.

In comparison, tedizolid is an oxazolidinone-class antibiotic that works by inhibiting mitochondrial protein translation [28, 106]. Inhibition of mitochondrial translation in venetoclax-resistant AML cells was observed to activate (or amplify) one or more branches of the ISR, which as we saw in Section 2, is a complicated pathway involving several distinct chemical species and regulatory mechanisms at both the RNA and protein levels. Thus, tedizolid would be expected to significantly affect the expression of several genes, including a subset of those that regulate c-Myc and Chop. Under these circumstances, especially with limited protein data, the mean-field approach is indispensable for capturing the underlying biological effects and offering interpretability to the experimental data.

In this work, perturbations to the regulatory gene network resulted from the administration of therapeutic agents. The time evolution of drug concentrations were modeled using a system of two coupled ordinary differential equations, with one equation describing the intra-cellular drug concentration, and the other the extra-cellular drug concentration. We showed that this simplistic approach captures the *in vivo* drug pharmacokinetics over several days by appropriately incorporating the drug half-life and the time to peak drug concentration into the model parameters. We also showed that for short-term *in vitro* experiments, taking the extra-cellular drug decay rate to be zero served as an excellent approximation to experimentally measured drug uptake curves. However, due to cellular metabolism and natural drug decay, the assumption of zero extra-cellular drug decay may not be appropriate for *in vitro* experiments that are conducted over a long period of time, such as several days, when there is no drug replenishment. Over these longer time intervals, the non-zero changes resulting from background drug decay may not be insignificant. However, due to the lack of *in vivo* metabolic and elimination processes in the *in vitro* settings, the rate of decay *in vitro* would be expected to be smaller than the *in vivo* drug decay rate. For these reasons, we modeled the extra-cellular *in vitro* drug decay to be non-zero, but smaller than the corresponding *in vivo* drug decay rate, during long-term *in vitro* experiments. As described in Section 4.2, this choice was supported by the non-monotonic behaviour of the protein-level data and the time-scale over which protein-level changes occurred.

The mean-field model predicted slow drug uptake by the molm-13 R2 cells, with the peak intracellular drug concentration achieved at approximately 36 hours after drug administration. While the quantitative result may be a reflection of the simplicity of the mathematical model, qualitatively, RNA sequencing data supported the notion that there is a biological basis for slow drug uptake by the resistant AML cells. In particular, differential gene expression analysis and gene ontology analysis indicated that, in comparison to drug sensitive molm-13 cells, the resistant cell line exhibited significant changes to the cell membrane structure and the formation of intra-cellular and extra-cellular vesicles, consistent with a reduced rate of drug uptake. Further analysis of the RNA sequencing data also indicated that the molm-13 R2 cells exhibited a significant decrease in CD38 and CD93 RNA levels in comparison to the parental drug-sensitive cell line, with the log_2_(fold change) = −4.12 and −6.67, respectively, and adjusted *p*-value = 0 for both changes. The change in CD38 expression was ranked as the second-most significant change and CD93 as the twelfth-most significant change by differential gene expression analysis with DEseq [98]. Interestingly, the CD38 and CD93 expression levels are important biomarkers for leukemia cell immunophenotype: CD34^+^CD38^−^ cells are a primitive sub-population of progenitor cells which are generally highly quiescent [107], while the cell surface lectin CD93 is expressed on a sub-population of these cells, identifying them as predominantly cycling, non-quiescent leukemia-initiating cells [108]. Therefore, the changes in CD38 and CD93 RNA expression exhibited by molm-13 R2 cells are consistent with a phenotypic shift toward a more stem-like quiescent cell. These observations were also reinforced by a 33% decrease in c-Myc RNA levels in the molm-13 drug-resistant cells compared to the sensitive cell line, *p*-value = 1.95×10^−59^.

These findings suggest that the mean-field approach can be considered a data-driven technique with the ability to infer deep biological insight that is not apparent from simple analysis of experimental data. For example, another prediction of the mean-field model was the existence of significant positive co-operativity for the stability effects of c-Myc with combination treatment, as well as for the production effects of Chop with combination treatment. Biologically, several distinct mechanisms can lead to cooperativity in the pathways that regulate c-Myc and Chop expression. Importantly, the mean-field model predicted the existence of co-operativity in the combined drug effects, despite not having knowledge of these individual mechanisms.

In the case of c-Myc, co-operativity might arise during protein stabilization and destabilization processes since there are several pathways that control the stability of the protein, and some of the stability mechanisms depend on prior phosphorylation by a subset of the pathways. For example, phosphorylation at Thr58 by GSK3*β*, which tags the protein for degradation, requires prior phosphorylation at Ser62 by ERK1/2, which increases the protein stability [31]. Furthermore, the PI3K/AKT pathway inhibits GSK3*β*, which provides a potential mechanism of co-operativity with ERK1/2 for increasing the overall stability of the c-Myc protein. The RNA sequencing data for resistant AML cells indicated that there was a slight upregulation in AKT, ERK1, and GSK3*β* during venetoclax treatment and/or combination therapy, see Figure 7. However, several other mechanisms that could co-operatively increase c-Myc stability, such as phosphorylation at Ser62 by SIRT1, PIM1 or ERK2, and phosphorylation at Ser329 by PIM2, were all significantly downregulated under each treatment condition, and particularly with combination treatment, see Figure 7. These findings are consistent with the predictions of the mean-field model.

In the case of Chop, co-operativity in production might arise due to the combination therapy activating or amplifying several distinct branches of the ISR. Indeed, it was hypothesized [27, 28] that both venetoclax and tedizolid individually lead to sub-lethal activation of the ISR, and that their combination leads to a heightened stress response that is sufficient to induce apoptosis in venetoclax-resistant AML cells, though the specific mechanism remains unclear. Experimental studies indicated [28] that venetoclax and tedizolid monotherapies both lead to structural changes that are consistent with mitochondrial stress in resistant AML cells. In other work, venetoclax treatment was observed to lead to increased ROS levels, particularly the production of peroxides [109], as well as an upregulation in mitochondrial transcription factor A (TFAM) and SOD2 protein levels, and several metabolic changes [105]. These changes could reflect a compensatory nuclear transcriptional response to mitochondrial (and perhaps low-level endoplasmic reticulum [110]) stress induced by treatment, enabling the cells to adapt and survive. The disruption of mitochondrial translation that results from tedizolid treatment could then serve to halt this adaptive mechanism and potentially lead to heightened stress and activation of additional stress response pathways. Importantly, due to the cross-talk between the mitochondria and endoplasmic reticulum (particularly via calcium signalling), stress in one of these membrane-bound organelles can trigger stress or dysfunction in the other [110–112], potentially activating another specific stress response or the more general ISR. Regardless of the specific mechanism, it is clear that cooperativity, as predicted by the mean-field model, plays a significant role in the combined drug effects that promote Chop expression.

The mean-field model also predicted that the weights *w_j,m,ℓ_* deviate from unity for the c-Myc production effects with combination treatment. This indicates that each drug does not contribute equally to the combination treatment, which could potentially be due to additional feedback mechanisms in the system that are activated during combination treatment, for example, due to heightened activation of the ISR. In particular, we see in Table S2 that the combination treatment weight for c-Myc and venetoclax treatment was near 0, while for tedizolid treatment it was above unity. Restricting the weight for tedizolid treatment to be equal to 1, it was possible to obtain a similar fit to the data with a slightly larger value of the weight for venetoclax treatment. This parameter flexibility is one drawback to the approach introduced here and illustrates why it is beneficial to validate the model predictions with an independent data set, such as RNA sequencing. In the absence of additional data, the pathways contributing most significantly to specific drug effects might not be individually identified; however, the relative contribution from each drug to the combined effect can at least be qualitatively estimated.

In closing, we remark that when building a mathematical model to describe a physical or biological process, one must consider the trade-off between the complexity of the model and the number of unknown parameters. When the experimental data set is small, there may not be enough data to uniquely parameterize the model, thus the focus should be to develop an approach that can capture the important overall interactions and relevant behaviour of the system. Here we developed such an approach which enables the identification of biological insights from a limited experimental data set. While we illustrated the power of this approach, we note that it does have important limitations that should be carefully addressed. For example, if the parameter ranges for fitting are chosen to be outside the correct biological ranges, the approach could potentially predict the incorrect interactions. Thus it is important to further validate the predictions of the model with additional experimental data if possible. In this case, we did this using RNA sequencing data, which provided a method to integrate RNA-level data with protein-level data. In a future work, we will investigate the changes to the Bcl-2 family protein levels that result from combination venetoclax and tedizolid therapy in venetoclax-resistant AML cells. We will use the method developed here to model the regulation of c-Myc and Chop expression, which both influence the commitment to cellular apoptosis.

## Acknowledgements

M.K. and M.P. acknowledge the financial support from the Canadian Institutes of Health Research (CIHR).

## Supplementary Information

### A Parameter values

**Table S1:**
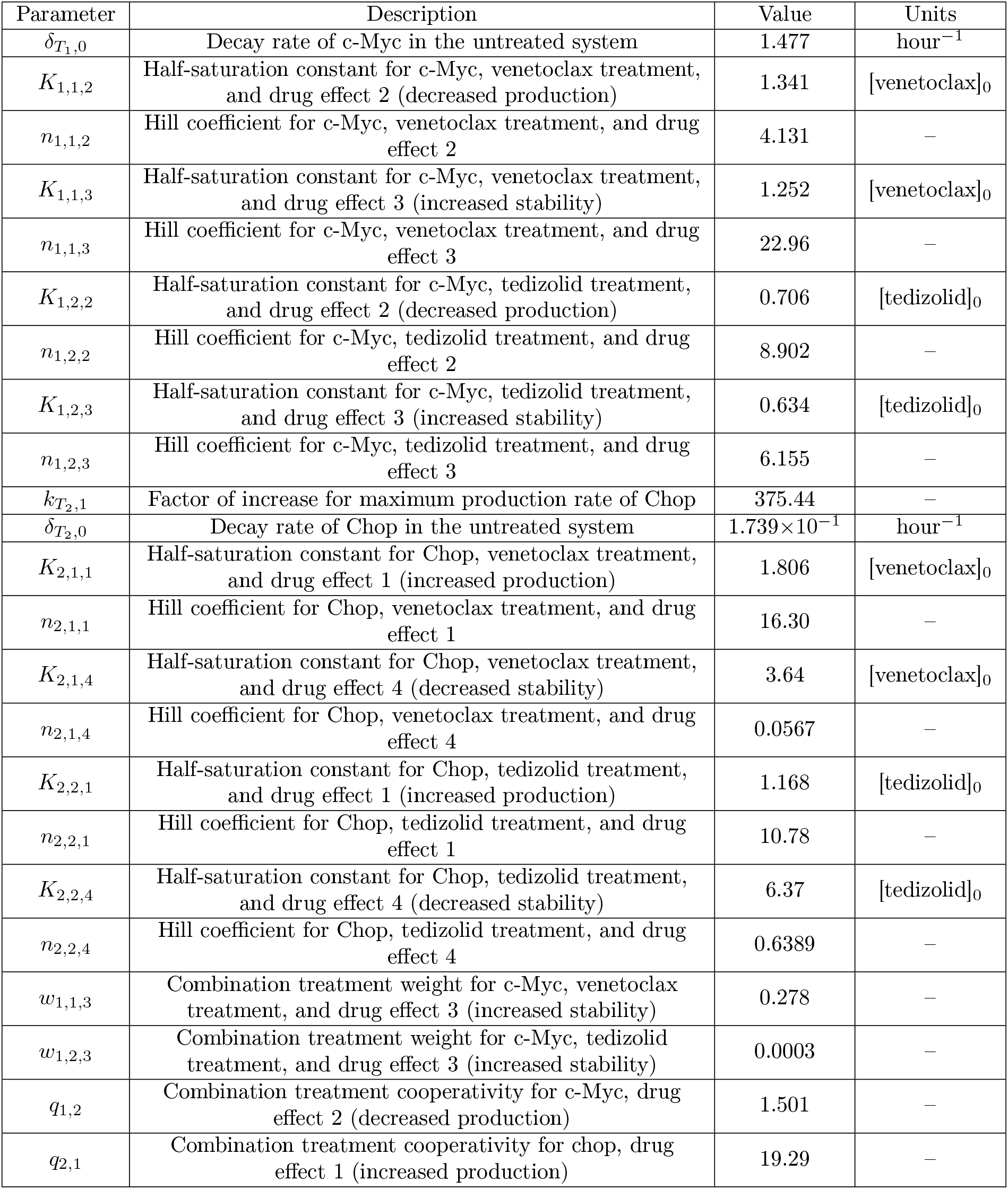
Nominal kinetic parameter set for the mean-field model, obtained from fitting to the experimental data.

| Parameter | Description | Value | Units |
| --- | --- | --- | --- |
| $\delta_{T_1,0}$ | Decay rate of c-Myc in the untreated system | 1.477 | hour<sup>-1</sup> |
| $K_{1,1,2}$ | Half-saturation constant for c-Myc, venetoclax treatment, and drug effect 2 (decreased production) | 1.341 | [venetoclax]<sub>0</sub> |
| $n_{1,1,2}$ | Hill coefficient for c-Myc, venetoclax treatment, and drug effect 2 | 4.131 | – |
| $K_{1,1,3}$ | Half-saturation constant for c-Myc, venetoclax treatment, and drug effect 3 (increased stability) | 1.252 | [venetoclax]<sub>0</sub> |
| $n_{1,1,3}$ | Hill coefficient for c-Myc, venetoclax treatment, and drug effect 3 | 22.96 | – |
| $K_{1,2,2}$ | Half-saturation constant for c-Myc, tedizolid treatment, and drug effect 2 (decreased production) | 0.706 | [tedizolid]<sub>0</sub> |
| $n_{1,2,2}$ | Hill coefficient for c-Myc, tedizolid treatment, and drug effect 2 | 8.902 | – |
| $K_{1,2,3}$ | Half-saturation constant for c-Myc, tedizolid treatment, and drug effect 3 (increased stability) | 0.634 | [tedizolid]<sub>0</sub> |
| $n_{1,2,3}$ | Hill coefficient for c-Myc, tedizolid treatment, and drug effect 3 | 6.155 | – |
| $k_{T_2,1}$ | Factor of increase for maximum production rate of Chop | 375.44 | – |
| $\delta_{T_2,0}$ | Decay rate of Chop in the untreated system | $1.739 \times 10^{-1}$ | hour<sup>-1</sup> |
| $K_{2,1,1}$ | Half-saturation constant for Chop, venetoclax treatment, and drug effect 1 (increased production) | 1.806 | [venetoclax]<sub>0</sub> |
| $n_{2,1,1}$ | Hill coefficient for Chop, venetoclax treatment, and drug effect 1 | 16.30 | – |
| $K_{2,1,4}$ | Half-saturation constant for Chop, venetoclax treatment, and drug effect 4 (decreased stability) | 3.64 | [venetoclax]<sub>0</sub> |
| $n_{2,1,4}$ | Hill coefficient for Chop, venetoclax treatment, and drug effect 4 | 0.0567 | – |
| $K_{2,2,1}$ | Half-saturation constant for Chop, tedizolid treatment, and drug effect 1 (increased production) | 1.168 | [tedizolid]<sub>0</sub> |
| $n_{2,2,1}$ | Hill coefficient for Chop, tedizolid treatment, and drug effect 1 | 10.78 | – |
| $K_{2,2,4}$ | Half-saturation constant for Chop, tedizolid treatment, and drug effect 4 (decreased stability) | 6.37 | [tedizolid]<sub>0</sub> |
| $n_{2,2,4}$ | Hill coefficient for Chop, tedizolid treatment, and drug effect 4 | 0.6389 | – |
| $w_{1,1,3}$ | Combination treatment weight for c-Myc, venetoclax treatment, and drug effect 3 (increased stability) | 0.278 | – |
| $w_{1,2,3}$ | Combination treatment weight for c-Myc, tedizolid treatment, and drug effect 3 (increased stability) | 0.0003 | – |
| $q_{1,2}$ | Combination treatment cooperativity for c-Myc, drug effect 2 (decreased production) | 1.501 | – |
| $q_{2,1}$ | Combination treatment cooperativity for chop, drug effect 1 (increased production) | 19.29 | – |

**Table S2:** Kinetic parameter set for the mean-field model with temporal delays in a subset of the drug effects, obtained by fitting to the experimental data.

| Parameter | Description | Value | Units |
| --- | --- | --- | --- |
| $\delta_{T_{E,1}}$ | Extra-cellular decay rate of venetoclax | $2.3 \times 10^{-3}$ | hour<sup>-1</sup> |
| $\delta_{T_{i,1}}$ | Intra-cellular decay rate of venetoclax | $9.7 \times 10^{-2}$ | hour<sup>-1</sup> |
| $\delta_{T_{E,2}}$ | Extra-cellular decay rate of Tedizolid | $8.9 \times 10^{-3}$ | hour<sup>-1</sup> |
| $\delta_{T_{i,2}}$ | Intra-cellular decay rate of Tedizolid | $8.1 \times 10^{-2}$ | hour<sup>-1</sup> |
| $k_{T1,1}$ | Factor of increase for maximum production rate of c-Myc | 11.43 | – |
| $\delta_{T1,0}$ | Decay rate of c-Myc in the untreated system | $3.851 \times 10^{-1}$ | hour<sup>-1</sup> |
| $K_{1,1,1}$ | Half-saturation constant for c-Myc, venetoclax treatment, and drug effect 1 (increased production) | 2.147 | [venetoclax]<sub>0</sub> |
| $n_{1,1,1}$ | Hill coefficient for c-Myc, venetoclax treatment, and drug effect 1 | 3.284 | – |
| $K_{1,1,4}$ | Half-saturation constant for c-Myc, venetoclax treatment, and drug effect 4 (decreased stability) | 3.703 | [venetoclax]<sub>0</sub> |
| $n_{1,1,4}$ | Hill coefficient for c-Myc, venetoclax treatment, and drug effect 4 | 1.683 | – |
| $K_{1,2,1}$ | Half-saturation constant for c-Myc, tedizolid treatment, and drug effect 1 | 4.849 | [tedizolid]<sub>0</sub> |
| $n_{1,2,1}$ | Hill coefficient for c-Myc, tedizolid treatment, and drug effect 1 | 2.670 | – |
| $K_{1,2,4}$ | Half-saturation constant for c-Myc, tedizolid treatment, and drug effect 4 (decreased stability) | 1.568 | [tedizolid]<sub>0</sub> |
| $n_{1,2,4}$ | Hill coefficient for c-Myc, tedizolid treatment, and drug effect 4 | 6.944 | – |
| $k_{T2,1}$ | Factor of increase for maximum production rate of Chop | 382.49 | – |
| $\delta_{T2,0}$ | Decay rate of Chop in the untreated system | $1.756 \times 10^{-1}$ | hour<sup>-1</sup> |
| $K_{2,1,1}$ | Half-saturation constant for Chop, venetoclax treatment, and drug effect 1 (increased production) | 1.288 | [venetoclax]<sub>0</sub> |
| $n_{2,1,1}$ | Hill coefficient for Chop, venetoclax treatment, and drug effect 1 | 20.23 | – |
| $K_{2,1,4}$ | Half-saturation constant for Chop, venetoclax treatment, and drug effect 4 (decreased stability) | 5.735 | [venetoclax]<sub>0</sub> |
| $n_{2,1,4}$ | Hill coefficient for Chop, venetoclax treatment, and drug effect 4 | 0.9208 | – |
| $K_{2,2,1}$ | Half-saturation constant for Chop, tedizolid treatment, and drug effect 1 (increased production) | 1.556 | [tedizolid]<sub>0</sub> |
| $n_{2,2,1}$ | Hill coefficient for Chop, tedizolid treatment, and drug effect 1 | 10.74 | – |
| $K_{2,2,4}$ | Half-saturation constant for Chop, tedizolid treatment, and drug effect 4 (decreased stability) | 8.772 | [tedizolid]<sub>0</sub> |
| $n_{2,2,4}$ | Hill coefficient for Chop, tedizolid treatment, and drug effect 4 | 0.6346 | – |
| $w_{1,1,1}$ | Combination treatment weight for c-Myc, venetoclax treatment, and drug effect 1 (increased production) | $1.039 \times 10^{-4}$ | – |
| $w_{1,2,1}$ | Combination treatment weight for c-Myc, tedizolid treatment, and drug effect 1 (increased production) | 2.423 | – |
| $q_{1,4}$ | Combination treatment cooperativity for c-Myc, drug effect 4 (decreased stability) | 208.90 | – |
| $q_{2,1}$ | Combination treatment cooperativity for chop, drug effect 1 (increased production) | 113.76 | – |
| $\tau_{1,1,1}$ | Time delay for c-Myc, venetoclax treatment, and drug effect 1 (increased production) | 10.36 | hour |
| $\tau_{1,2,1}$ | Time delay for c-Myc, tedizolid treatment, and drug effect 1 (increased production) | 27.42 | hour |
| $\tau_{2,2,1}$ | Time delay for Chop, tedizolid treatment, and drug effect 1 (increased production) | 5.028 | hour |
| $\tau_{2,2,4}$ | Time delay for Chop, tedizolid treatment, and drug effect 4 (decreased stability) | 6.00 | hour |

